# Age distinguishes selection from causation in cancer genomes

**DOI:** 10.1101/2025.10.06.680730

**Authors:** David Cheek, Martin Blohmer, Martin Nowak, Tibor Antal, Kamila Naxerova

## Abstract

Cancer genome sequencing efforts have revealed hundreds of genes under positive selection, many of which are now being developed as therapeutic targets. However, positively selected mutations also populate our aging tissues in the absence of cancer. For most mutations, it is currently unknown whether they are recurrently found in cancer genomes because they cause cancer or because they expand during normal tissue evolution and are passively inherited. Here, we develop a mathematical and statistical framework that distinguishes these two factors. We discover – across thousands of cancer and normal tissue genomes – that mutations that most strongly increase cancer risk are enriched in younger patients’ cancers, whereas mutations that are positively selected in normal tissue without causing cancer are enriched in older patients. Focusing on a particularly data-rich cancer type, acute myeloid leukemia, we show that genetic differences between young- and adult-onset cancers can largely be explained by the cumulative effects of normal tissue evolution, contradicting the long-standing notion that childhood cancers require a distinct set of causal mutations. Our framework establishes patient age as a powerful resource for clarifying whether positively selected mutations in cancer genomes are truly disease-promoting.

## MAIN

Which mutations cause cancer? To understand the mechanisms of carcinogenesis and towards the development of treatments, this question continues to occupy cancer biology after decades of research. The answers so far have primarily been derived from cancer genomes viewed through a Darwinian lens. Cancer is conceptualized as an extreme outcome of natural selection among somatic cells, where one group of cells comes to proliferate excessively at the expense of the whole. If a mutation is found in cancer genomes more commonly than expected under a model of neutral evolution, then the mutation is thought to be under positive selection during carcinogenesis and is flagged as potentially causal for cancer development^1–9^.

However, a wealth of sequencing studies published within the last 10 years have shown that positive selection is ubiquitous in normal tissues^10–15^, calling into question whether all positively selected mutations increase the risk of malignant transformation. In fact, emerging evidence suggests that many variants that are expanding in normal tissue do not contribute to cancer causation. For example, multiple genes that are under positive selection in liver disease rarely appear in hepatocellular carcinoma^11,13^. Similarly, among esophageal squamous cell carcinomas (ESSCs), non-synonymous mutations in the *NOTCH1* gene are far more common than synonymous mutations, indicating strong positive selection and therefore implicating *NOTCH1* mutations in ESSC development by standard methods; yet refuting that implication, *NOTCH1* mutations are even more frequent in the normal esophageal epithelium than in ESSCs, suggesting that the signals of positive selection in ESSC genomes are an inheritance from normal tissue evolution, and more provocatively that the mutations might inhibit carcinogenesis^16,17^. Mouse experiments agree, demonstrating that *Notch1* mutations can drive clonal expansion in the normal esophageal epithelium but impair tumor growth^18^.

It may seem paradoxical at first glance that any mutation could drive clonal expansion and not cause cancer, but in many scenarios, expansion and carcinogenesis might be non-overlapping. For example, a mutation could increase a stem cell’s probability of symmetric self-renewal but have no effect on its absolute proliferation rate or invasive properties, as has been shown for *Notch1*^18^. However, standard methods of cancer driver identification based on signals of positive selection in cancer genomes do not delineate carcinogenesis from decades of normal tissue evolution that may have preceded it.

Comparisons of cancer and normal tissue genomes have provided hope for the statistical disentanglement of selection and causation, and here we aim to establish a rigorous inference framework based on such data. For most cancer types though, normal tissue studies remain limited in terms of cohort sizes. Causal information’s availability in cancer genomes without a normal tissue control therefore requires further interrogation. For this, we take a population genetics view of the timeline of somatic evolution predating cancer. We reason that because growth of mutant clones in normal tissues takes time, signals of positive selection in cancer genomes are inherited from normal tissue in increasing magnitudes towards older ages. On the other hand, we propose that the most potent cancer-causing mutations expedite carcinogenesis, thereby appearing preferentially in younger patients’ cancers. We regard patient age data alongside cancer genomes as a potential reservoir of untapped information on selection and causation.

## RESULTS

### Defining a mutation’s carcinogenic effect

To assess a specific mutation’s contribution to cancer development while allowing ignorance of other causal variables, we took a standard statistical view of causation. We defined a mutation’s carcinogenic effect by its influence on a cell’s probability of initiating a malignant growth that is eventually diagnosed. Consider two hypothetical cells, statistically identical in every way except that one cell has mutation z while the other does not. The cell with mutation z initiates cancer at some rate r_1_ (probability per unit time) while the other cell initiates cancer at rate r_0_. Then the per cell hazard ratio for cancer initiation given the mutation, r_1_/r_0_, defines the mutation’s carcinogenic effect (Figure 1A). A mutation’s effect on a tissue’s rate of cancer is determined by the mutation’s carcinogenic effect and its frequency in the tissue (Figure 1B).

**Figure 1:**
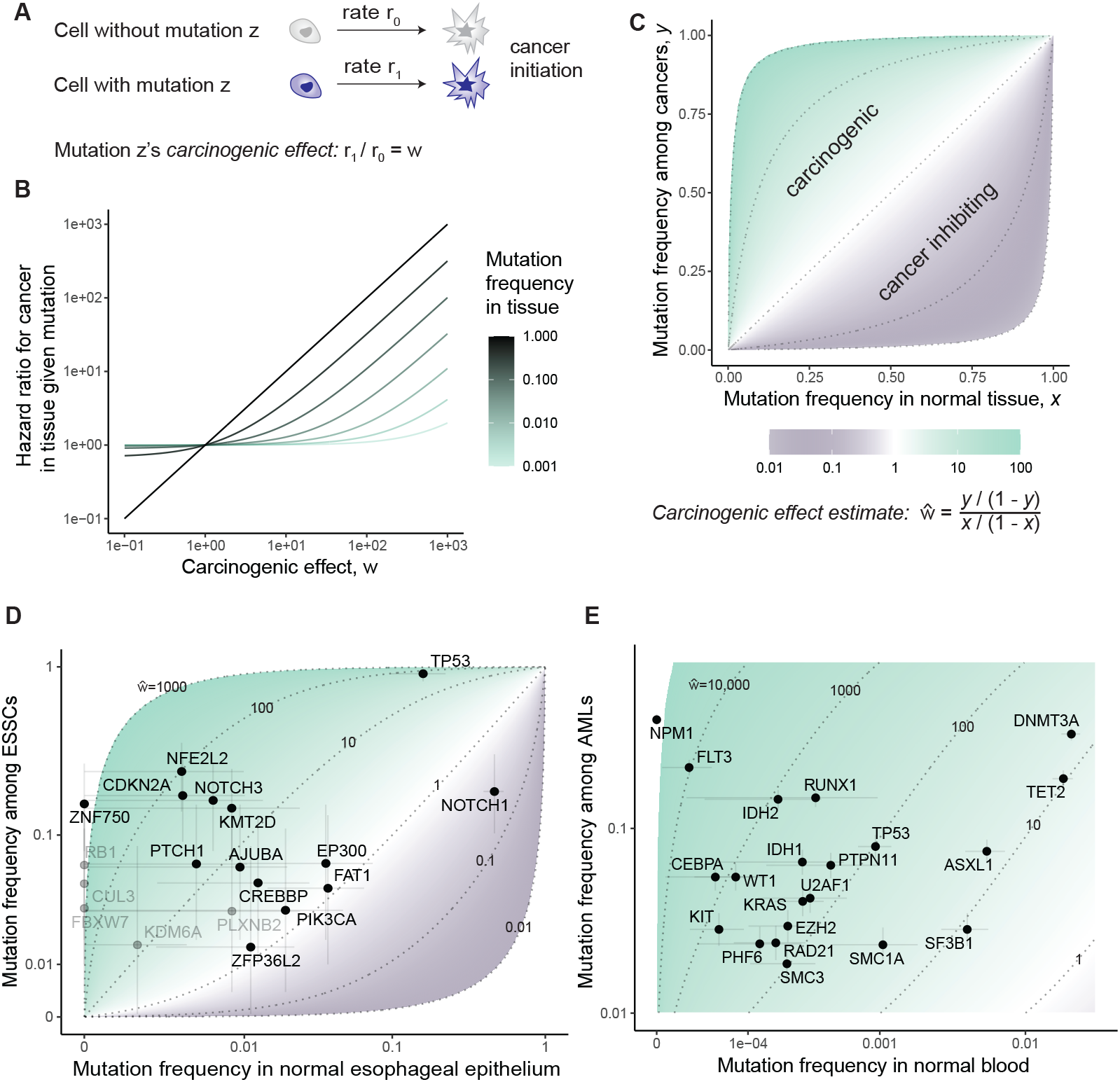
Mutation frequencies in normal tissue and cancer genomes estimate carcinogenic effects. (A) Carcinogenic effect definition. Cell with mutation z initiates cancer at rate r_1_. Cell without mutation z, otherwise statistically identical, initiates cancer at rate r_0_. Mutation z’s carcinogenic effect is r_1_/r_0_. (B) Relationship between mutation’s carcinogenic effect (x-axis), fraction of cells with mutation in tissue (color), and cancer rate in tissue given mutation’s presence normalized by cancer rate without mutation (y-axis). (C) Relationship between mutation’s carcinogenic effect (light green - carcinogenic; grey - protective), fraction of normal cells with mutation among many people (x-axis), fraction of cancer founder cells with mutation (y-axis). (D,E) Carcinogenic effects estimated per gene for esophageal squamous cell carcinoma (ESSC, D) and acute myeloid leukemia (AML, E). (D) Fraction of 63 ESSC samples with non-synonymous mutations or small indels in each gene (y-axis; with jitter for visualization) against estimated fraction of cells mutated in patient-matched normal esophageal epithelium samples (x-axis). Data from Yokoyama et al.^17^ Genes exhibiting positive selection in ESSC^17^ are plotted. Genes mutated in fewer than 5 samples are light colored. (E) Fraction of 2340 AML samples^41–43^ with non-synonymous mutations or small indels in each gene (y-axis) against estimated fraction of cells mutated in 2410 age-matched normal blood samples^39,40^. AML driver genes from Bailey et al.^44^ are plotted. (D,E) Horizontal and vertical error bars respectively represent 95% bootstrap and binomial confidence intervals for mutation frequencies. Dotted lines indicate carcinogenic effect estimates.

The carcinogenic effect can also be thought of in terms of an imaginary randomized controlled trial where normal tissue cells from many people are randomly assigned or disallowed mutation z, and some of the cells subsequently found cancer. Say that fraction *x* of the normal tissue cells is assigned the mutation and that fraction *y* of cancer founders carry the mutation. We derived (Methods) that the odds ratio

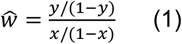

of mutation frequencies in cancer relative to normal tissue estimates the mutation’s carcinogenic effect (Figure 1C): greater enrichment of the mutation in cancer-founder relative to normal cells means greater carcinogenicity.

### Mutation frequencies in cancer and normal tissue estimate carcinogenic effects

Mutations can of course never be artificially induced in humans to test causality, and therefore the carcinogenic effect by our definition can never be perfectly measured. Nevertheless, the hypothetical experiment that defines carcinogenicity has a measurable natural analogue provided by the genetic heterogeneity of human tissues, where some cells carry mutation z while others do not. The frequency *x* of mutation z among normal tissue cells can be estimated from deep sequencing of normal tissue samples, meanwhile the frequency *y* of mutation z among cancer founder cells can be estimated by the fraction of cancers where mutation z is clonal, enabling calculation of the odds ratio (1).

At this point, it is important to emphasize that somatic evolution does not necessarily distribute the mutation among cells independently of other variables that might influence carcinogenesis. For example, an epistatic interaction between two mutations in normal tissue could lead to the two mutations preferentially appearing together in normal tissue cells. Or mutational processes could act heterogeneously across cells in a tissue (as has been described for APOBEC mutagenesis^19^). More generally, if a mutation’s presence in a normal tissue cell is associated with another variable that causes cancer (such as another mutation), then the mutation’s odds ratio in cancer relative to normal tissue (1) could over- or under-estimate the carcinogenic effect, depending on whether the association is positive or negative. With this caveat in mind, we appealed to the odds ratio as a rough, practical estimate for the carcinogenic effect. For a first view of carcinogenic effect estimates in human data, we turned to the esophagus, a tissue whose somatic evolution is relatively well documented.

Yokoyama et al^17^ performed targeted sequencing of purported driver genes among 68 esophageal squamous cell carcinoma (ESSC) samples and 68 physiologically normal esophagus samples from the same 68 patients. To compare malignant and non-malignant cells, we wanted to ensure that the samples of apparently normal tissue do not represent clones from which the cancers arose (Theory Supplementary Note), hence, we removed from the analysis 5 patients who exhibited a common mutation between cancer and normal tissue samples. Then for each gene, focusing on non-synonymous point mutations and indels, we compared the fraction of ESSCs mutated against the fraction of normal epithelial cells mutated (mutant cell fraction calculations detailed in Methods) to estimate carcinogenic effects per gene. We do not distinguish carcinogenic effects of different mutations within a gene, which would be possible with larger datasets. The resulting panorama provides a unique view onto esophageal carcinogenesis, showcasing a spectrum of cancer-causing potency among genes (Figure 1D). *TP53* and *NFE2L2* emerge as the main genes implicated in the development of ESSC, with carcinogenic effect estimates standing in the hundreds. This comes as no surprise in the case of *TP53*, which is arguably the best-understood cancer-associated gene. Ample evidence from mouse models proves that *TP53* loss causes cancer in a variety of tissues, both in the germline^20,21^ and in somatic cells^22–25^. Activating alterations in *NFE2L2*, coding for the stress response transcription factor NRF2, have not yet been shown to cause cancer in mice, partially because mice with activated NRF2 signaling in the germline die at young ages due to esophageal hyperkeratosis^26^. However, its outstanding mutational enrichment in cancer relative to normal tissue genomes, which far exceeds other, more frequently mutated genes in ESSC, should motivate future efforts to clarify its carcinogenicity in animal models. The other end of the spectrum is more notable, however. Several well-known cancer-associated genes (e.g. *EP300, FAT1*) have carcinogenic effect estimates in the statistical vicinity of one, yet they also exhibit strong signals of positive selection in ESSCs^17^. These statistics suggest that cancer causation and positive selection in the esophagus are not equivalent – which has been previously implied^16,17^, although without estimates of carcinogenic effects. One of the best understood examples of this non-equivalence is *NOTCH1*, the second most mutated gene in ESSC, whose mutations we infer to inhibit ESSC initiation – consistent with prior discussions^16^ and experimental investigation^18^ showing that *Notch1* loss impairs tumorigenesis in the mouse esophageal epithelium. Another thought-provoking data point is *PIK3CA*, a gene that shows strong signals of positive selection in ESSC genomes and is widely regarded as a major cancer driver gene across many tissue types^27–29^, but here inferred to contribute relatively little to ESSC initiation – consistent with a recent mouse study showing only a modest carcinogenic effect for *Pik3ca* mutations^30^.

Next, we turned to the blood. Mutations under selection in normal hematopoietic stem cells have been studied extensively^31–38^, making the blood a particularly well-illuminated trial ground for our framework. Here, we used high depth (approximately 1000x) targeted sequencing of AML-associated genes among 2410 peripheral blood samples from Abelson et al.^39^ and Fabre et al.^40^. We compared mutation frequencies in the normal blood samples to 2340 acute myeloid leukemia (AML) samples, comprising 466 samples from COSMIC^41^, 633 samples from Bottomly et al.^42^, and 1874 samples from an NCRI study^43^ that have the same age range as the normal blood samples (36 to 103 years). We focused on non-synonymous variants in 21 genes that were sequenced in all normal blood and AML studies and that are part of a standard list of AML driver genes^44^. As in the esophagus, so in the blood, we observed a wide span of carcinogenic effects (Figure 1E). We estimated carcinogenic effects around 10 for the most common clonal hematopoiesis driver genes *ASXL1, DNMT3A, TET2*, and *SF3B1*, and a carcinogenic effect of ∼100 for *TP53*. Meanwhile mutations in *FLT3, CEBPA, IDH2*, and *WT1* increase a cell’s transformation rate by about 800- to 10000-fold. The most frequently mutated gene in AML yet unmutated in the normal blood data is *NPM1*, indicating an exceptionally powerful carcinogenic effect. Broadly consistent with our results, people with clonal hematopoiesis have an increased risk of future AML: the risk increase is modest for mutations in *ASXL1, DNMT3A*, and *TET2* and greater for *SF3B1, TP53*, and *IDH1/2*^39,45,46^, though for genes uncommonly mutated in normal blood, associations with future AML lack statistical resolution.

We note that carcinogenic effect estimates are overall higher for the blood than the esophagus. One conceivable explanation for this difference is that copy number alterations, rather than SNVs and small indels, are more important for esophageal carcinogenesis. It could also be that a greater number of mutations is required for – and therefore each mutation contributes less to – ESSC than AML. However, there is a technical difference between the cancer types too in this analysis: the normal esophagus samples are patient-matched to the cancer samples, whereas the normal blood samples are from people with no history of blood malignancies. The blood’s unmatched comparison is more exposed to confounding variables. People who develop AML may statistically differ from those who don’t with respect to multiple variables, such as smoking, diet, or germline genetics, which may influence both the rate of mutant clonal expansions in the blood and a person’s propensity to develop AML^47–49^, potentially augmenting some mutations’ frequencies in AML relative to normal blood samples beyond what would be explained by the mutations’ carcinogenic effects alone.

We are not concerned about mutagenesis as a confounder for our analyses of ESSC and AML, because these cancers have mutation burdens similar to single cell-derived colonies of the respective normal tissues^17,50^ (Extended Data Figures 1A,B), but other tissues might not exhibit such balanced distributions of mutation numbers between cells. Relatedly, some mutations arriving during cancer growth may appear at clonal variant allele frequencies in a sample, exaggerating the mutation burden of cancer relative to normal tissue. Among 2411 colorectal cancers in COSMIC, the mean exome-wide mutation burden is 348, whereas among 1387 normal colonic crypts^14,51^, the mean exome-wide mutation burden is only 21 (Extended Data Figure 1C). To compare colorectal cancer and the normal colon in terms of specific purported driver mutations while reducing biases by mutagenesis pre- or post-carcinogenesis, we focused on 718 colorectal cancer samples (from COSMIC^52^) whose mutation burden is in the same range as the normal colonic crypts, and we employed a logistic regression model with mutation burden and age as covariates to estimate carcinogenic effects among driver genes (Methods; Extended Data Figures 1D,E). Again, we measured a large range for mutations’ effects on the per cell probability of carcinogenesis. We estimated that *TP53, APC, KRAS*, and *PIK3CA* mutations have carcinogenic effects in the hundreds. Genes such as *FBXW7* and *PTEN* appear one order of magnitude less important, while *ARID1A* has a carcinogenic effect that does not significantly differ from one. We acknowledge that mutations’ effects could differ in the context of hypermutated cancers, which are not included in our analysis.

For the future, ever richer catalogues of mutations in normal tissues, especially patient-matched to cancers, will enable increasingly well-controlled and high-resolution quantifications of mutations’ effects both on clonal expansion in normal tissue and on cells’ risk of cancerous transformation. However, the need to determine cancer causation is immediate. For now, cancer genetic data are copious while normal tissue data remain comparatively limited. We therefore endeavored to disentangle causation and selection from cancer genomes without a normal tissue control.

### Selection in normal tissue and carcinogenicity differentially shape mutations’ age distributions in cancers

Is it possible to extract causal information from cancer genomes alone? In the remainder of the manuscript, we will develop and test a framework that enables partial identification of causal genes without access to mutation frequencies from the cancer’s normal tissue of origin. Our proposed approach leverages a data type that has not played a dominant role in the interpretation of cancer genomes so far: patient age. The idea is twofold. Firstly, in normal somatic evolution, older tissues have experienced a longer duration of selective pressures than younger tissues. Hence, due to mitotic inheritance from normal tissue to cancer, selection in normal tissue tends to have a greater impact on cancer genomes of older patients. Secondly, we reasoned that the more powerful cancer-causing mutations should more vigorously accelerate the onset of carcinogenesis, therefore appearing in relatively young patients’ cancer genomes. We thus hypothesized that patient age data may contribute insights into somatic mutations’ selective and carcinogenic effects that are inaccessible from cancer genomes alone.

To develop the idea formally, we looked to express in mathematical terms the relationship between mutation frequencies in cancer genomes, patient age, and parameters of lifelong somatic evolution. For a simple first model, we began with a branching process to illustrate the stochastic arrival of a mutation and the selection that acts upon it in a normal tissue (Figures 2A and 2B). The model’s starting point is a tissue at birth that is composed of N cells, all without some mutation z. Then each cell in the tissue without mutation z either divides at rate a_0_, is lost through death or differentiation at rate b_0_ (with a_0_=b_0_), or acquires mutation z at rate v. On the other hand, each cell with mutation z divides or is lost at respective rates a_1_ and b_1_. So the mutation may edit the division, death or differentiation rates of a cell. The composite parameter s=a_1_-b_1_ is the mutation’s selective effect in normal tissue.

**Figure 2:**
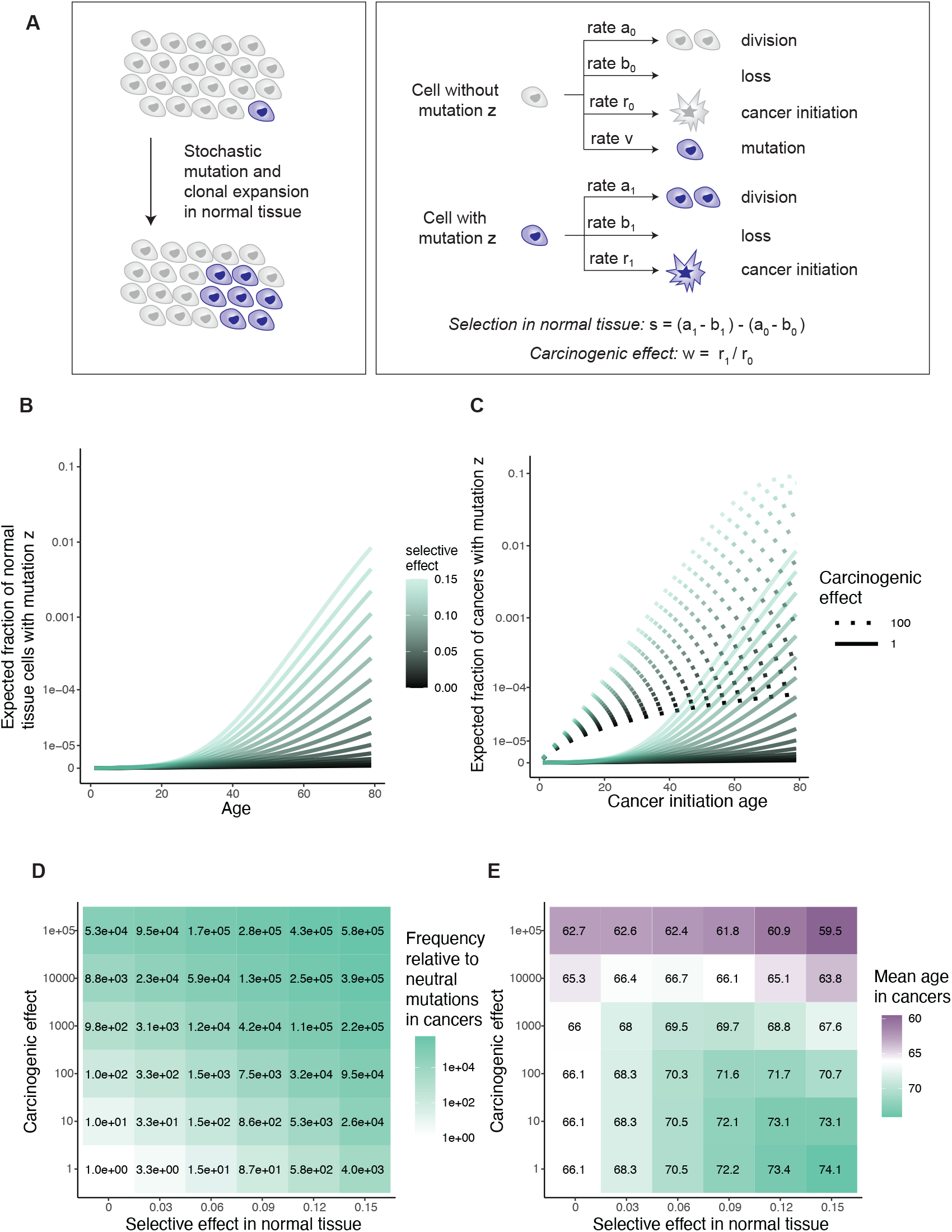
Selection and carcinogenicity differentially shape mutations’ age distributions in cancers. (A) Branching process model of a normal tissue’s somatic evolution illustrates stochastic cell division, loss (through death or differentiation), acquisition of mutation z, and cancer initiation. Difference between growth rate of cells with and without mutation z is the selective effect in normal tissue. Ratio of rates at which cells with and without mutation z initiate cancer is the carcinogenic effect. (B,C) Theoretical mutation frequency among normal tissue cells (B) and cancer-initiating cells (C) as a function of mutation’s selective effect in normal tissue, carcinogenic effect, and tissue age. (D) Selection in normal tissue and cancer causation lead to similar mutation frequencies in cancer genomes. Colors represent frequency of cancer with mutation z divided by frequency of cancer with neutral mutation of same acquisition rate. (E) Selection in normal tissue and cancer causation lead to differing age distributions for mutations in cancer genomes. Colors represent mean patient age of cancers carrying mutation z. (B–E) parameter values: lifespan 80 years; stem cell pool size N=10^5^; mutation rate per year v=10^−8^; division and loss rate a_0_=b_0_=1 per year without mutation and a_1_=1+s, b_1_=1 with mutation; baseline carcinogenesis rate r_0_=10^u^t^3^, where u is uniformly distributed on (−16,-13), caricaturing that cancer risk rises with age and varies between people.

Versions of this model have been studied for decades in population genetics^53^, and recently it has been applied to somatic blood evolution^15,47,54^, with theoretical predictions largely consistent with thousands of single-time-point blood samples^15,54^ as well as hundreds of longitudinal blood samples^40^. Thus, the model can be regarded as a reasonable description of mutation and selection in the blood, while for other tissues whose somatic evolution is not yet so well understood, the model can more prudently be considered a cartoon. We extended the evolutionary model described so far to additionally encompass carcinogenesis: cells transform to cancer at some rate, the rate possibly altered by the presence of mutation z (Figure 2A). The central question then is how the selective and carcinogenic effect parameters manifest in cancer genetic data.

We calculated mutation z’s expected frequency among normal tissue cells (Figures 2C) and among cancer-initiating cells (Figure 2D, Extended Data Figure 2A), as a function of tissue age and the model parameters. To check consistency with our original framework (Figure 1C), we compared the mutation’s frequency between cancer and normal tissue via the odds ratio stratified by age, recovering the carcinogenic effect (Methods). Then to explore the hypothetical absence of normal tissue data, we calculated the odds ratio for mutation z’s presence in cancer genomes relative to a neutral mutation with the same mutation rate, which resembles dN/dS. This statistic has a positive relationship with both the selective effect in normal tissue and with the carcinogenic effect (Figure 2D, Extended Data Figure 2A; Theory Supplementary Note), illustrating that dN/dS alone cannot distinguish selection from causation.

Next, we sought to assess a mutation’s preference towards younger or older cancer genomes. For this, a straightforward summary statistic is the mean age of patients whose cancers carry mutation z. We found that in the setting of no cancer causation (carcinogenic effect = 1), the selective effect in normal tissue has a positive relationship with the mean age. By contrast for any specific selective effect, the carcinogenic effect has a negative relationship with the mean age, representing that carcinogenic mutations speed up carcinogenesis (Figures 2E, Theory Supplementary Note). Although the precise numerical details of these relationships are sensitive to other somatic evolutionary parameters, which may vary greatly between tissues, the directionality of the relationships is robust (Extended Data Figure 2B).

Thus, our simple model laid out so far predicts differences between mutations in their patient age distributions, due to mutation-specific selective and carcinogenic effects. We note a complication to the model that could contribute additionally to age-dependent cancer genetics: it could be that mutations’ carcinogenic effects are themselves age dependent. Age-dependent effects might arise for example due to mutational interactions with the aging epigenetic state of stem cells, or with other somatic mutations who accumulate with age. Relatedly, young cancer patients might be more likely to have cancer-predisposing germline mutations, potentially interacting with specific somatic mutations. Overall, there are several possible reasons to expect that driver mutation frequency spectra differ between older and younger cancer patients. Careful empirical investigation is therefore required to assess whether age distributions may help to distinguish selection and causation in real-world cancer data.

Hence, we wanted to check four questions in data. (I) Do patient age distributions differentiate mutations? (II) Can age differences between mutations be explained by age-dependent carcinogenic effects? Ideally, the answers to (I) and (II) would be positive and negative respectively, inviting the next two questions. (III) Are powerful carcinogenic mutations found preferentially in younger patients’ cancers? (IV) Are mutations who are positively selected in normal tissue found preferentially in older patients’ cancers? We begin with question (I).

### Cancer types differ in the degree of age divergence between driver genes

To quantify the degree of age divergence between purported driver genes per cancer type in COSMIC^52^, we calculated the variance between purported driver genes (using an established list of driver genes^44^) of patient ages at which non-synonymous SNVs/small indels appear, normalized by the overall variance of the patient age distribution. Comparing cancer types, we observed a range of degrees of age divergence between driver genes, with esophageal and colorectal cancers exhibiting relatively little and acute myeloid leukemia exhibiting the most (Figure 3A). Indeed, it is well known that the genetic makeup of childhood and adult AMLs differs starkly^55^. For a high-powered view of driver genes’ age dependence in AML, we collated four datasets (the three datasets that contributed to our analysis of adult AML in Figure 1E, now including all patient ages, and a pediatric AML dataset^55^) totaling 4736 AML patients from ages 0 to 100 (Extended Data Figure 3A). We calculated the mean patient age for non-synonymous SNVs/small indels per driver gene, observing wide separation of genes (Figure 3B). For example, *KIT* has mean age 36 while *TET2* has mean age 63. Given driver genes’ age differences, and since normal somatic evolution has been documented far more comprehensively in the blood than in other organs, we decided that AML would be the ideal primary exploration ground for the somatic evolutionary meaning of age distributions posed by questions (II), (III), and (IV).

**Figure 3:**
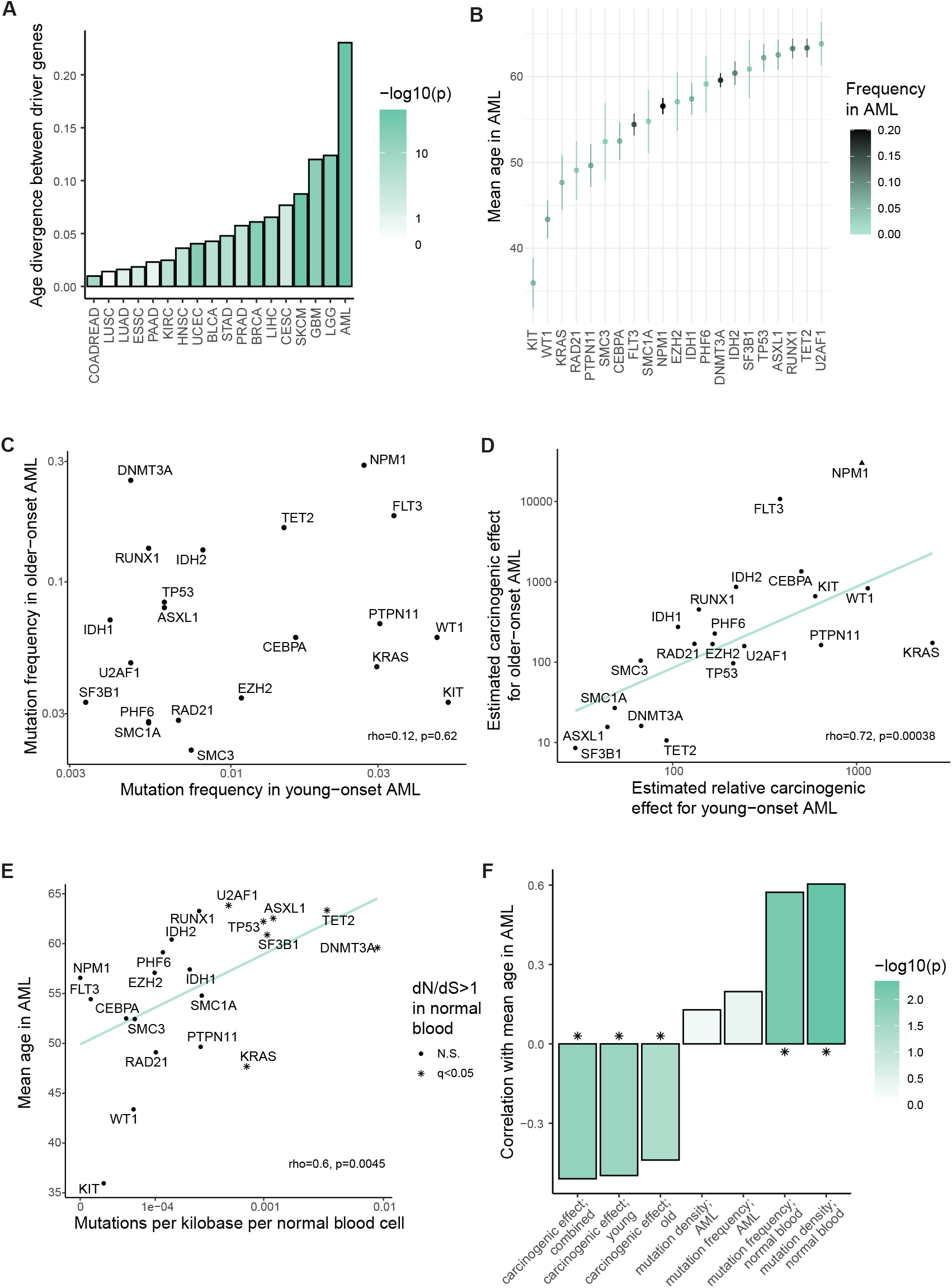
Age distributions of AML mutations provide information on selection and carcinogenicity. (A) Among cancer types, AML exhibits greatest age divergence between driver genes. Age divergence statistic is defined by variance between driver genes of non-synonymous SNVs/small indels’ patient age, normalized by overall variance of patient age per cancer type (Methods). Cancer type-specific driver genes listed in Bailey et al.^44^. p-values (color) from Kruskal-Wallis test of patient age vs. driver gene. Data from COSMIC (Methods). See Supplementary Table 3 for unabbreviated cancer types. (B) Mean age of AML patients for non-synonymous SNVs/small indels per driver gene (Methods). Error bars are 95% confidence intervals. Four AML datasets collated (Extended Data Figure 3A). (C) Fraction of older-onset AML (mean age 61, standard deviation 12) vs. fraction of younger-onset AML (mean age 11, standard deviation 7) samples with non-synonymous SNV/small indel per gene. rho=0.12, p=0.62. (D) Carcinogenic effect estimates for older AML (Figure 1D) vs. relative carcinogenic effect estimates for younger AML (normalized by median carcinogenic effects in older AML, Extended Data Figure 3); rho=0.72, p=0.00038. *NPM1* has infinite carcinogenic effect estimate for older-AML, included at finite value for visualization. Line represents regression excluding *NPM1*. (E) Mean AML patient age for gene’s mutations vs. number of mutations per gene per kilobase per normal blood cell (rho=0.6, p=0.0045). Mutation densities in normal blood estimated from 2410 samples^39,40^; donor age range 36 to 103. Genes under positive selection in normal blood according to dN/dS>1 with q<0.05^40^ are denoted by star shape; these genes have older mean AML patient ages (Wilcoxon test, p=0.016). (F) Mutation frequencies in normal blood (normalized and unnormalized by gene length; data as in E) and carcinogenic effect estimates (younger- and older-AML, and combined score (see Extended Data Figure 3E)) have distinct relationships with AML patient age, according to Spearman’s correlation among driver genes. Mutation frequencies in AML (normalized and unnormalized by gene length) do not have significant correlation with AML patient age.

### Carcinogenic effects largely align between young- and older-onset AML

To develop an explanatory picture for AML genetics’ age dependence, first addressing question (II), we wanted to estimate carcinogenic effects for young-onset AML, enabling comparison with our estimates for older-adult AML in Figure 1E. For a young-onset AML cohort, we selected patients younger than 25 years old from the collated AML datasets (Extended Data Figure 3A), totaling 1481 patients whose age distribution has mean 11 and standard deviation 7; by contrast, our adult-onset AML cohort comprises 2973 patients whose age distribution has mean 61 and standard deviation 12. Among AML driver genes, we found a large discrepancy between mutation frequencies among childhood and adult AML patients (Figure 3C), which is in line with previous observations^55^ and is commonly interpreted as diverging biology between the two AML types^56^. For example, mutations in *DNMT3A* and *RUNX1* are common in later-onset AML but uncommon in childhood AML, while *KIT* is the only gene whose mutations are more common in young AML patients. Overall, for this subset of AML-associated genes, no correlation between mutation frequencies in the younger and older patients could be detected (rho=0.12, p=0.62).

While our carcinogenic effect estimates for adult AML were calculated by the odds ratio of mutation frequencies in adult AML relative to age-matched normal blood (Figure 1E), we wanted to similarly estimate carcinogenic effects in childhood AML. The new difficulty for childhood AML inference is that mutations in normal childhood blood are rarely observed, with variant allele frequencies too low for reliable detection^36^, which incidentally is in alignment with the theory that selection in normal tissue has had less cumulative effect on younger people’s tissues. Towards inference in the absence of observable mutations, we approximated a gene’s mutation frequency in normal childhood blood by the gene’s rate of mutation acquisition which we in turn approximated as proportional to the gene’s length (Methods). These two approximations amount to the simplifying assumptions of a uniform mutation rate along the genome and that selection has negligible impact on mutation frequencies in normal childhood blood. Thus, we obtained a benchmark against which to judge mutation frequencies in childhood AML, leading to estimates of relative carcinogenic effects (Extended Data Figure 3B). Now, childhood and adult AML carcinogenic effect estimates can be compared.

In contrast to the disagreement of mutation frequencies between old and young AML (Figure 3C), we found that carcinogenic effect estimates strongly correlate between the old and young age groups (rho=0.72, p=0.00034; Figure 3D). That is, while the ranking of mutation frequencies is radically revised with the progression of age, the ranking of carcinogenic effect estimates is more stable – a result that is robust to changes in the age boundaries that define the patient groups (Extended Data Figure 3C). Nevertheless, some specific genes do exhibit substantial changes in their estimated carcinogenic effect ranking from childhood to adulthood. Employing bootstrap resampling of AML and normal blood samples, at least 95% of bootstraps support that that *FLT3* has a carcinogenic effect ranked more than three places higher in adult AML compared to childhood AML and that *KRAS* has a carcinogenic effect ranked more than six places higher in childhood relative to adult AML (methods; Extended Data Figure 3D). However, we note a caveat of these apparent changes with age, that differences between childhood and adult carcinogenic effect estimates could be exaggerated by the (necessarily) distinct methodological origins of the childhood and adult normal blood references (Methods). In any case, the differences between childhood and adult AML in terms of carcinogenic effect estimates (Figure 3D) are far more limited than our pre-theory expectation based only on mutation frequencies in AML (Figure 3C), indicating that age-dependence of carcinogenic effects is unlikely to be the primary explanatory factor underlying the age-dependence of AML mutation frequencies.

### Carcinogenic effect estimates in AML negatively associate with mutations’ patient age

We next looked to test whether those mutations found towards younger patients’ AML genomes tend to be the more powerful cancer-causing mutations (question (III)), which theory predicts to be so (Figure 2E). For a summary statistic of age, we again used the mean patient age of mutations per driver gene (Figure 3B). Among genes, we observed that mean age in AML negatively associates with carcinogenic effect estimate (rho=-0.44, p=0.048 for adult-onset AML carcinogenic effect estimates; rho=-0.5, p=0.023 for young-onset AML carcinogenic effect estimates; Extended Data Figure 3E). This negative relationship is consistent with the concept that causation corresponds to a quickening of carcinogenesis. According to our framework, the other explanatory factor that may underlie mutations’ age distributions in AML is selection in normal blood.

### Mutation frequencies in normal blood positively associate with AML patient age

To compare genes in terms of their degree of selection in normal blood (question (IV)), we calculated for each gene the number of non-synonymous mutations per kilobase per normal blood cell (Methods), based on the same normal blood data^39,40^ as for the adult AML carcinogenic effect estimates (Figure 1E). We observed that across genes, their mutation density in normal blood strongly correlates with mean age in AML (rho=0.61, p=0.0038; Figure 3E). For another measure of selection, we turned to dndscv^8^, a highly parametrized version of dN/dS that accounts for mutation rate heterogeneity along the genome. In Fabre et al.^40^ among 1593 normal blood samples, positive selection was evidenced for 7 out of our 21 AML-associated genes (dN/dS>1 with q<0.05), and we observed that these positively selected genes tend to have older mean ages in AML (Wilcoxon test, p=0.016; Figure 3E). In summary, positive selection in normal blood manifests preferentially in AML genomes of older patients, which stands contrary to the inverse relationship between carcinogenic effect estimates and mean ages in AML (Figure 3F). Thus, for the best-documented tissue of the body, causation of cancer and selection in normal tissue leave distinct signals in patient age distributions.

### Young-age biases of driver mutations in cancers indicate carcinogenicity

Having evidenced that mutations’ age distributions carry information on selection and carcinogenicity for AML, we endeavored to expand our investigation to other cancer types where normal tissue data are lacking, to carefully extract what causal information may be available in age distributions. Without extensive normal tissue sequencing data, we did not wish to impose any single specific model of somatic evolution, and so, more humbly than parameter inference, we intended to identify genes whose mutations are carcinogenic. This requires a falsifiable prediction of the null hypothesis of no carcinogenicity. The null hypothesis states that while a mutation may or may not be positively selected in normal tissue, the mutation has no influence on the rate of transformation to cancer. More formally, we defined the null hypothesis as a collection of models of somatic evolution, encompassing our original branching process model (Figures 2A, 4A) in addition to models where clonal expansions in normal tissue follow sub-exponential growth curves – including Gompertz, logistic, and polynomial models (Extended Data Figure 4A), which may be more realistic for solid tissues due to tissue architecture or clonal interference^57^ – with the unifying restriction that the carcinogenic effect of the mutation under consideration is fixed at one throughout life (see Theory Supplementary Note for precise scope of model class).

We found that one simple prediction holds universally across the space of null models: for a mutation that is positively selected in normal tissue but does not influence cancer, the mutation is found in cancer genomes of older patients relative to neutral mutations (Figures 2E, 4B, Extended Data Figure 4B, Theory Supplementary Note). This prediction is universal because it follows from the definition of positive selection: in normal tissue, a positively selected mutation on average grows in frequency relative to neutral mutations as age progresses, and if this mutation does not influence carcinogenesis, then its old-age bias would be bequeathed to cancer genomes. Crucially, the old-age bias for positively selected mutations predicted by the null hypothesis can be demonstrated false. If a mutation both (a) exhibits positive selection in cancer genomes (by having a greater frequency than the background neutral mutation rate) AND (b) is enriched towards *younger* patient ages relative to neutral mutations, then this would constitute evidence to reject the null hypothesis (Figure 4C), in favor of the alternative hypothesis that the mutation is carcinogenic.

**Figure 4:**
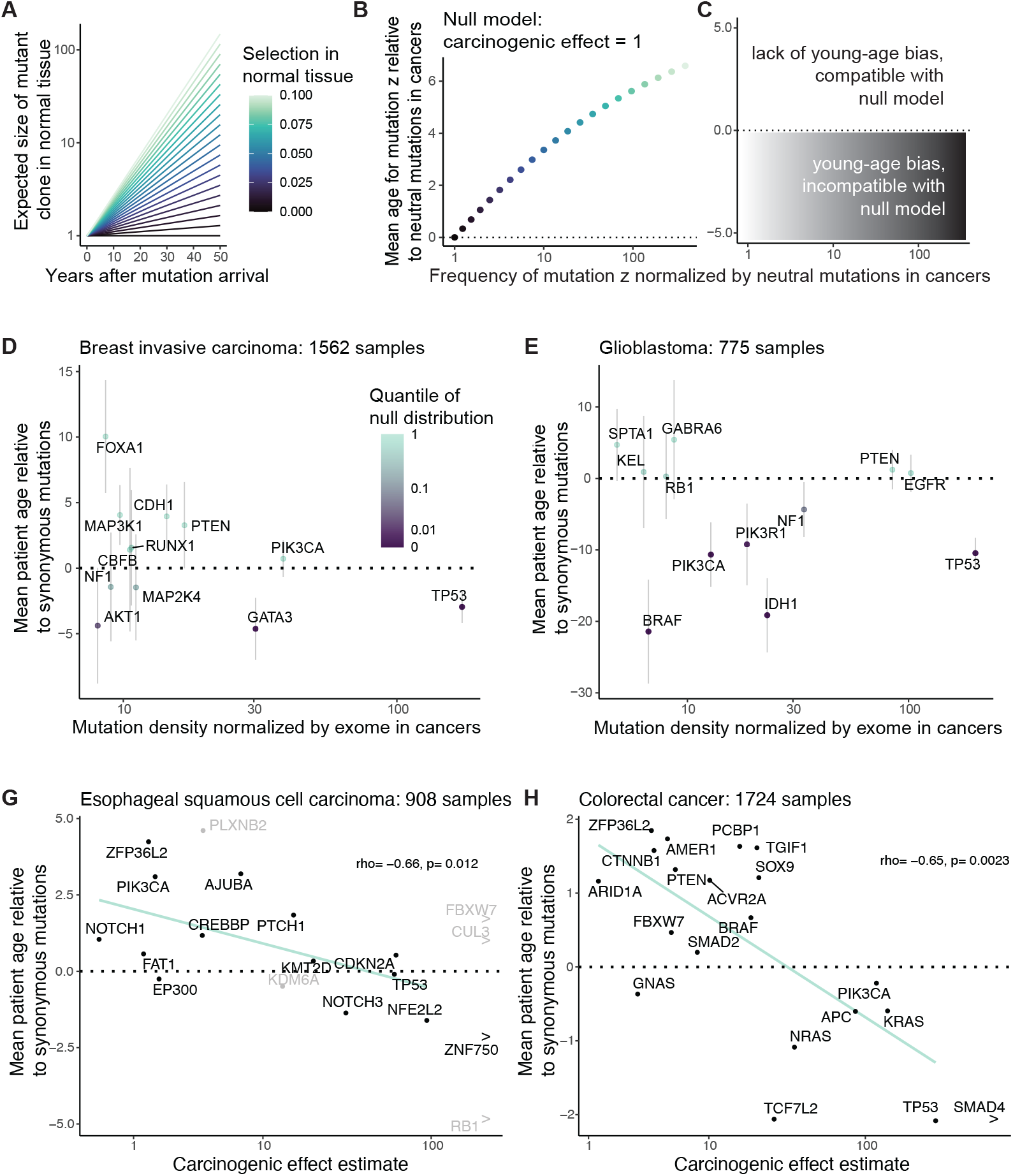
Young age biases in cancer genomes indicate carcinogenicity. (A,B) Mutation frequency and age statistics according to branching process model. (A) Expected growth of positively selected mutant clones in normal tissue. (B) Positively selected non-carcinogenic mutations are found preferentially in older patients’ cancers relative to neutral mutations. Y-axis shows the mean patient age for cancers with mutation z minus the mean patient age for neutral mutations. X-axis shows expected frequency of mutation z in cancers divided by expected frequency of neutral mutation with the same acquisition rate. (A,B) Color denotes mutation z’s selective effect in normal tissue. Generalization in Extended Data Figures 4A,B and Theory Supplementary Note. (C) Null hypothesis test schematic. Mutations with elevated frequency AND young-age bias in cancer genomes relative to neutral mutations are incompatible with null model of no carcinogenicity. (D,E) Null hypothesis test for driver genes in breast invasive carcinoma (BRCA;D) and glioblastoma (GBM; E). 1562 BRCA samples and 775 GBM samples from COSMIC. Y-axis shows mean patient age for non-synonymous mutations and small indels per driver gene minus the mean patient age for synonymous mutations in exome (Methods; for short, *age difference*). Error bars are 95% Wald confidence intervals. Color denotes quantile of age difference statistic with respect to null distribution, generated by permuting patient ages (Methods). X-axis is number of mutations per gene divided by number of mutations in exome across the cohort, normalized by coding sequence length. Genes are from Bailey et al.^44^; top 12 genes with greatest number of mutations per kilobase plotted. (F,G) Mean patient age per driver gene negatively associates with carcinogenic effect estimate in esophageal (rho=-0.66, p=0.012) and colorectal cancers (rho=-0.65, p=0.0023). Y-axis shows mean age per driver gene relative to synonymous mutations, as in (D,E). Patient ages from 908 ESSC samples and 1724 COADREAD samples in COSMIC. COADREAD samples with mutation burden greater than 251 excluded (to match mutation burden range of normal colon crypts; Methods). Carcinogenic effect estimates from Figure 1D and Extended Data Figure 1E. Genes with infinite carcinogenic effect estimates are denoted by “>“. Genes whose carcinogenic effect estimates are based on fewer than 5 samples are colored gray and excluded from correlation statistics and regression lines.

Before applying this null hypothesis test to cancer types for whom normal tissue data are lacking, we first applied the test to AML, focusing on the same 21 AML driver genes analyzed so far. Because these driver genes definitionally exhibit positive selection in AML samples, it only remained for us to test their mutations’ age distributions against neutral mutations. For this, we focused on samples with whole exome sequencing in COSMIC, so that exome-wide synonymous mutation could be used as the neutral benchmark. We removed samples with greater than ten times the median mutation burden, to avoid that an extreme minority of samples have an outsized effect on the synonymous mutation age distribution. Among the remaining 939 AML samples, we calculated the mean patient age weighted by the number of non-synonymous mutations per driver gene minus the mean patient age weighted by the number of synonymous mutations over the exome (Extended Data Figure 4C). For each gene, we assessed this age metric against a null distribution generated by permuting patient ages, and we observed that 6 out of the 21 genes have a significant young-age bias (q<0.05). These young-age biased genes, consistent with our earlier analyses of AML age distributions (Figure 3), tend to have fewer mutations in normal blood (although lacking statistical significance, Extended Data Figure 4D) and have significantly greater carcinogenic effect estimates overall (Extended Data Figure 4E). In conclusion, young age biases in AML can identify a subset of causal genes, and this subset tends to represent the more powerful carcinogenic effects.

We applied the null hypothesis test similarly to other cancer types in COSMIC (Supplementary Table 1, Extended Data Figure 4F), grouping cancers by their stem cell lineage of origin. Here, we stratified by stem cell lineage rather than molecularly defined cancer subtype because we are concerned with cancer genomes’ inheritance from decades of normal tissue evolution; it is within stem cells that mutation acquisition and selection have life-long consequences on the genetic landscapes of our tissues. For example, we amalgamated different subtypes of breast invasive carcinoma (BRCA), since they are all thought to descend from the mammary stem cell lineage^58^. Among 1562 BRCA samples, focusing on 23 established driver genes for BRCA (listed in Bailey et al.^44^), we found that *GATA3* and *TP53* mutations are biased towards young ages relative to exome-wide synonymous mutations (q<0.05; Figure 4D), inconsistent with unbiased inheritance from normal tissue evolution. The young-age bias for *TP53* mutations adds to the long-established evidence – including germline genetics^59,60^ and mouse studies^22,61^ – for *TP53* contribution to breast cancer risk. On the other hand, *PIK3CA* mutations, which are overwhelmingly prevalent in breast cancer, are not biased towards young ages. Analogously, among 775 glioblastoma patients in COSMIC, non-synonymous mutations in *TP53, IDH1, PIK3CA, BRAF*, and *ATRX* are significantly young-age biased relative to exome-wide synonymous mutations, but this is not so for *PTEN* and *EGFR* (Figure 4E). It must be reiterated that lack of evidence to reject the null hypothesis does not demonstrate lack of carcinogenicity, rather it says that further evidence beyond the cancer genetic and age data is required.

Among 908 ESSC samples in COSMIC, we found a significant young-age bias for *RB1*, evidencing causality (Supplementary Table 1). Indeed, *RB1* is classically and unambiguously implicated as a tumor suppressor gene for other cancer types by germline evidence. Concomitantly, among the 63 patient-matched ESSC and normal esophagus samples that led to carcinogenic effect estimates in Figure 1D, *RB1* mutations were found in 4 cancers and zero normal tissue samples, supporting high carcinogenic effects though more data are needed. Other genes did not exhibit statistically significant young age biases in ESSC, which could be because positive selection in the normal esophagus raises their age distribution (Figures 2C,E), or it could be that the data are underpowered to reject the null hypothesis for individual genes. Nevertheless, age distributions of the genes compared against each other provide information on carcinogenicity. We observed a negative association between ESSC driver genes’ mean ages and carcinogenic effect estimates (Figure 4G; rho=-0.66, p=0.012).

Among 2040 colorectal cancer samples in COSMIC, we found young age-biases for *TP53, TCF7L2*, and *SMAD4* with p<0.05, although only *TP53* remains statistically significant after correcting for multiple hypothesis testing (Extended Data Figure 4G, Supplementary Table 1). We moreover observed that mean ages negatively correlate with carcinogenic effect estimates, though not statistically significant (Extended Data Figure 4G, rho=-0.38, p=0.1). To reduce the possibility of biases introduced by mutagenesis (as suggested in Extended Data Figure 1C), we then recalculated mean ages exclusively among the 1724 COADREAD samples with exonic mutation burden less than 251, to match the mutation burden range of the normal colon crypt samples^14,51^ who we used for the carcinogenic effect estimates. With these low mutation burden COADREAD samples, we found that the negative correlation between mean ages and carcinogenic effect estimates strengthens to rho=-0.65 with p=0.0023 (Figure 4H). In summary, the theorized negative relationship between age and carcinogenicity (Figure 2F) held true for AML, ESSC, and COADREAD (cancer types with normal tissue data that enabled carcinogenic effect estimation).

### Chromosomal alteration age biases in AML associate with frequencies in normal blood

Having focused on SNVs/small indels so far, we next turned to somatic copy number alterations (SCNAs). SCNAs are a hallmark of cancer genomes^62^ and they are thought to exert significant carcinogenic effects, which we would like information upon. It is however unclear how well our framework can be applied to SCNAs. Due to technical difficulties of detecting SCNAs at low cell fractions in bulk samples of polyclonal normal tissue, the rules of somatic evolution are far less understood for SCNAs than SNVs/small indels, and the rules may take a different form. It is thought that SCNA acquisition is non-clocklike and sometimes explosive^63–66^, with a flood of copy number alterations late during carcinogenesis potentially overwriting the historical record of normal tissue evolution and shrouding the relationship between alterations’ carcinogenic effects and age. We therefore decided to investigate relatively simple cancer genomes with few SCNAs for information on selection and cancer causation.

For a case study cancer type with few chromosomal abnormalities and with uniquely voluminous normal tissue data, we looked again to AML. We examined SCNA frequencies in 2113 AML samples from Tazi et al.^43^ and 482,789 normal blood samples from the UK Biobank (SCNA calls by Loh et al.^67^). Previous estimates indicate that alterations present in at least ∼4% of cells per normal blood sample can be detected in the latter study^54^. We focused initially upon gains and losses of autosomal chromosome arms and whole chromosomes, since these alterations are available in both AML and normal blood datasets. As expected, alterations are far more numerous in AML: 32% of AMLs and 1% of normal blood samples have at least one SCNA; 14% of AMLs and 0.15% of normal blood samples have at least two SCNAs (Extended Data Figure 5A). To compare frequencies of specific alterations between AML and normal blood while ameliorating confoundment by genome instability, we restricted attention to the 1809 AML samples and the 482,043 normal blood samples with one or zero SCNAs. We observed that the most prevalent alteration in AML is amplification of chromosome 8; this alteration is observed in 5% of AMLs and has mean cell fraction 1.8e-4 in normal blood, with a carcinogenic effect estimate ∼1800 (Figure 5A). On the other hand, the most common alteration in normal blood is deletion of 13q, whose carcinogenic effect does not significantly differ from one. These results indicate that SCNAs – just like SNVs – have a wide range of carcinogenic effects.

**Figure 5:**
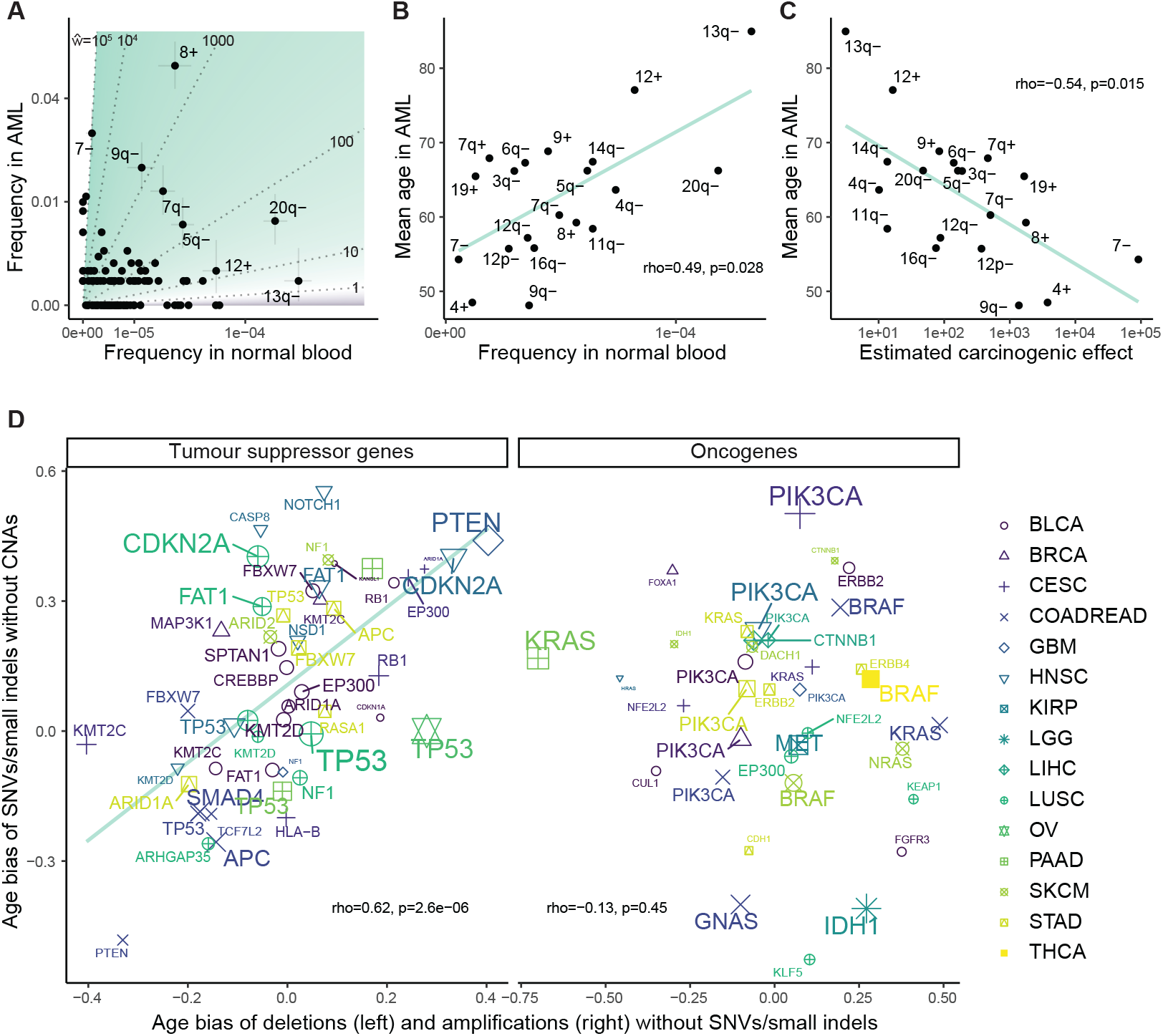
Selection and carcinogenicity shape age distributions of somatic copy number alterations. (A) SCNA frequencies in AML and normal blood estimate carcinogenic effects. Fraction of 1809 AML samples with each SCNA from Tazi et al.^43^ plotted against mean cell fraction estimate among 482,043 normal blood samples from the UK Biobank^67^ (normal blood donors aged 40-75). Samples with more than one SCNA excluded. Square-root scale. Vertical and horizontal error bars respectively represent 95% binomial and bootstrap intervals, shown for common SCNAs. (B) Mean age of AML patients with SCNA positively associates with mean cell fraction estimate of SCNA in normal blood (rho=0.49, p=0.028). Top 20 most frequent alterations over the normal blood and AML cohorts are plotted (alterations ranked by frequency in each cohort independently; we plot alterations with highest mean of the two rankings). (C) Mean age of AML patients with SCNA negatively associates with carcinogenic effect estimate (rho=-0.54, p=0.015). (D) Pan cancer comparison of age biases between SCNAs and SNVs/small indels. Left: For each cancer type and tumor suppressor gene, age bias of non-synonymous SNVs/small indels (rank biserial correlation between ages of patients with vs. without mutation in gene) among samples *without* SCNA is compared to the age bias of deletions among samples *without* non-synonymous SNV/small indels in that gene. rho=0.62, p=2.6×10^−6^. Right: For each cancer type and oncogene, age bias of non-synonymous SNVs/small indels among samples *without* SCNA is compared to age bias of amplifications among samples *without* non-synonymous SNV/small indels. rho=-0.13, p=0.45. See Supplementary Table 3 for unabbreviated cancer types. Driver genes are from Bailey et al.^44^. Genes with an SCNA and a SNV/small indel in at least 5% of samples per cancer type are plotted.

Next, we asked whether information on selection and carcinogenicity may be provided by SCNAs’ age distributions in AML. We calculated the mean age of AML patients with each amplification and deletion. Across the top 20 most common alterations in AML and normal blood, we observed a positive correlation between mean age in AML vs. frequency in normal blood (rho=0.49, p=0.028; Figure 5B), and a negative correlation between mean age in AML vs. carcinogenic effect estimate (rho=-0.54, p=0.015; Figure 5C), agreeing with our theory. These correlations are robust to the number of alterations under consideration (Extended Data Figure 5B). We note however that the correlations no longer hold if we include the minority of samples with many SCNAs (Extended Data Figure 5B), indicating that genome instability can weaken signals of selection and carcinogenicity.

### Distinct mutational disruptions of tumor suppressor genes share common age biases

For cancer types beyond AML, interpreting SCNA age distributions becomes more challenging. The difficulty is firstly that genome instability is more pervasive and secondly that normal tissue data are more limited. For a pan-cancer assessment of whether our framework applies to SCNA age distributions, in the absence of normal tissue controls, we appealed to SNVs/small indels in cancer genomes as a benchmark. We sought to compare age distributions of SCNAs against age distributions of SNVs/small indels that might be expected to exert similar phenotypic effects. The prime candidates for alterations with plausibly similar phenotypic effects are deletions of tumor suppressor genes (TSGs) vs. loss-of-function mutations in TSGs, as both alteration types ultimately abrogate gene function^4^. Other candidates are amplifications of oncogenes vs. gain-of-function variants in oncogenes. Demonstrating a deep relationship between these mutational types, Davoli et al.^4^ defined TSGs and oncogenes by their characteristic patterns of SNVs/small indels in cancers, showing that the numbers of TSGs and oncogenes per chromosome arm along with the strengths of signals of positive selection in those genes can predict the frequencies of chromosome arm losses and gains pooled across cancer types. These associations may be partially driven by co-occurrence of these alterations in the same cancer genome (i.e. loss-of-function mutation of one allele and deletion of the other, or gain-of-function of one allele and subsequent amplification of the mutant form)^68^, and conceivably, the associations might also be driven by phenotypic similarities between the mutational types without the need for co-occurrences. The latter scenario, if true, would enable age distributions of SNVs/small indels to serve as a suitable benchmark to assess age distributions of SCNAs. We reasoned that agreement between SNVs/small indels vs. SCNAs on their age distributions would support that our age distribution theory applies to SCNAs, as well as supporting the hypothesis that the distinct mutational types exert comparable carcinogenic and selective effects.

To put alterations (SCNAs or gene mutations) of different cancer types on the same scale, we measured an alteration’s age bias per cancer type by the rank-biserial correlation between patient age and alteration status (the effect size for a Mann-Whitney U test comparing the age distribution between patients with vs. without the alteration), which ranges from negative one to positive one, respectively representing bias towards younger or older patients. We applied this metric to alterations in the TCGA database (SCNAs called by Van Loo et al.^69^). To reduce the influence of genome instability and mutagenesis, we focused on samples without whole genome doubling. Also, we removed uterine corpus endometrial carcinoma and lung adenocarcinoma from the analysis, because they show significant negative correlations between patient age and number of synonymous mutations (respectively r=-0.15, p=5×10^−4^ and r=-0.13, p=0.3×10^−3^), in violation of our model’s prediction that neutral mutations accumulate with age. For each of the remaining cancer types, we focused on autosomal driver genes (again from the comprehensive driver gene list of Bailey et al^44^.). We calculated the age bias for non-synonymous SNVs/small indels per gene among samples <u>without</u> any SCNA in that gene.

We additionally calculated the age bias for each deletion and amplification per gene among samples <u>without</u> any non-synonymous SNV/small indel in that gene (Supplementary Table 2). The reason for avoiding cooccurrence of the mutational types is to compare the alterations’ separate effects without the complication of interactions.

Firstly, we focused on TSGs. We observed that age biases of deletions vs. SNVs/small indels in TSGs have a strong positive relationship: Spearman’s correlation is 0.62 (p=2.6e-6) among those TSG-cancer type pairs for which each alteration is present in at least 5% of samples (Figure 5C). This age agreement is robust to the specified lower bound of alteration frequencies per gene (Extended Data Figure 5C). We also employed a linear regression to assess the relationship between deletion vs. SNV/small indel age bias while accounting for cancer type as a covariate, still observing a strong positive association (Extended Data Figure 5D). Next, we turned to oncogenes. We observed that the age biases of amplifications vs. SNVs/small indels in oncogenes do not have a correlation significantly different from zero (r=-0.13, p=0.45; Figure 5D). The lack of age alignment between amplifications and mutations in oncogenes holds regardless of filtering parameters (Extended Data Figure 5B), and a linear regression that accounts for cancer type does not show a significant association between age biases of the two mutational types (Extended Data Figure 5D).

The age comparisons of SCNAs vs. SNVs/small indels are helpful on several counts. First, that the distinct mutational disruptions of TSGs agree on age distributions supports the notion that age distributions of SNCAs provide information on selection and cancer causation. Second, the comparisons support that deletions and mutations within TSGs exert broadly similar effects whereas amplifications and mutations within oncogenes overall have more divergent effects. Indeed, some mutations within oncogenes idiosyncratically alter protein structure and function in a way that amplifications cannot. Third, young-age biases provide orthogonal lines of evidence supporting the carcinogenicity of disrupting some TSGs (Supplementary Table 2).

### Multi-hit carcinogenesis: carcinogenic effects negatively associate with age and driver burden

Finally, we wanted to more explicitly express our framework in terms of the prevailing paradigm of cancer genetics, that carcinogenesis is the result of an array of alterations acting in concert. For a minimalist multi-mutation mathematical model that allows a variety of carcinogenic effects, we assumed that mutations arise randomly along each lineage in normal tissue at constant rate (as a Poisson process), and that each mutation has a carcinogenic effect that is independently sampled from a Gamma distribution; a lineage’s probability of cancerous transformation per unit of time is some baseline rate of carcinogenesis multiplied by the product of the mutations’ carcinogenic effects (Figure 6A).

**Figure 6:**
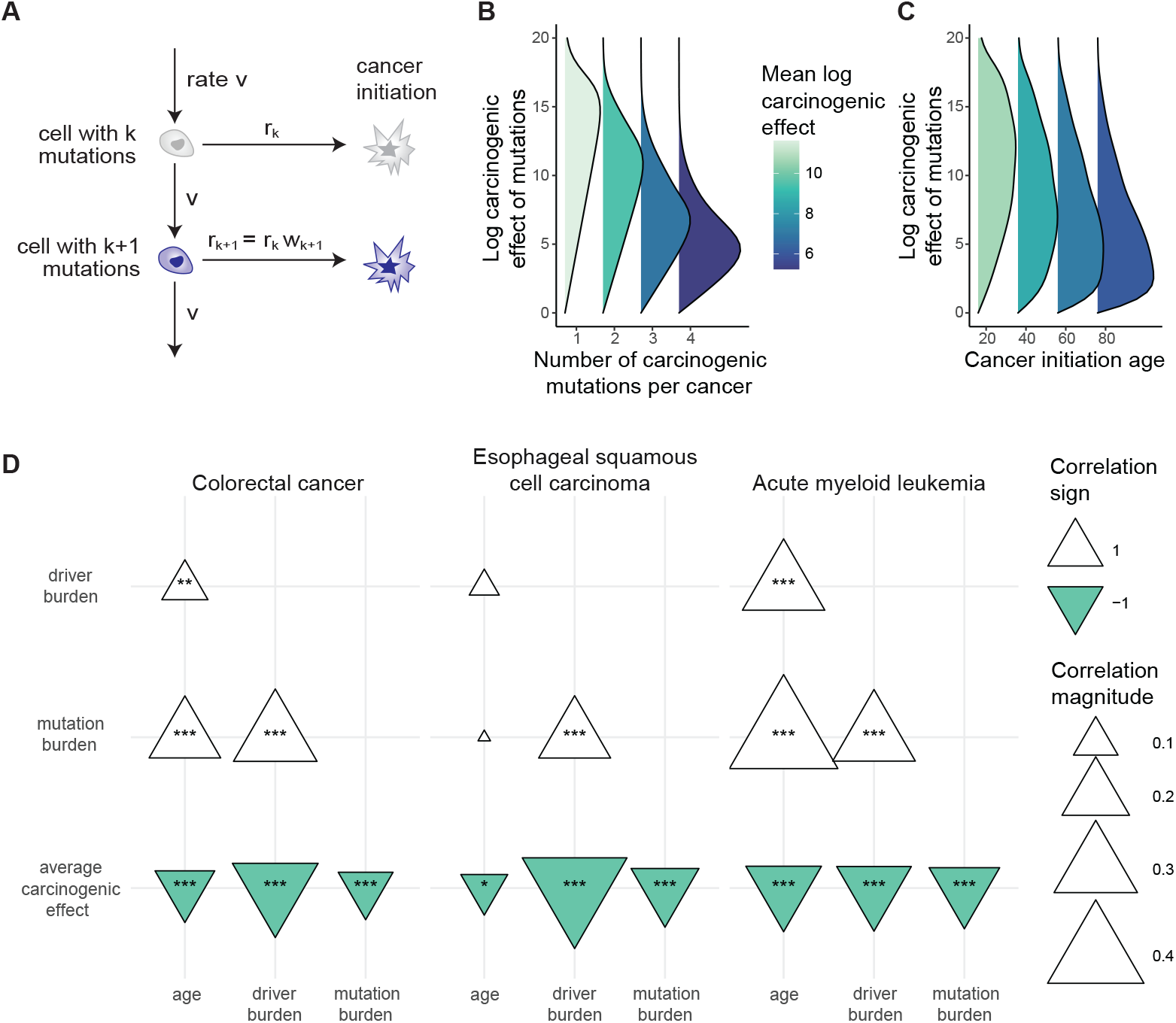
Multi-hit carcinogenesis: carcinogenic effects negatively correlate with age and driver mutation burden. (A) Schematic for model of multi-hit carcinogenesis. A lineage acquires carcinogenic mutations as a rate v Poisson process. Each new mutation has carcinogenic effect sampled from a distribution. The lineage’s rate of cancerous transformation is proportional to the product of the mutations’ carcinogenic effects. (B,C) Illustrational calculations: (B) Cancers with fewer carcinogenic mutations tend to have mutations with greater carcinogenic effects. (C) Cancers arising at younger ages tend to have mutations with greater carcinogenic effects. (B,C) parameters: lineage’s baseline rate of cancerous transformation without mutations is 10^−9^; mutation acquisition rate v=0.005 along lineage; mutations’ carcinogenic effects independently follow Gamma distribution with shape parameter 2 and rate parameter 1. (D) Multi-hit model predictions are qualitatively consistent with data for cancers with available carcinogenic effect estimates (COADREAD, ESSC, AML). Spearman correlations among cancer samples between patient age, mutation burden (number of exonic SNVs/small indels), driver burden (number of non-synonymous SNVs/small indels in purported driver genes), and mean rank carcinogenic effect of driver mutations per sample (for each cancer type, driver genes’ carcinogenic effects are ranked according to estimates in Figures 3C, 1D, and Extended Data Figure 1E). ∗, ∗∗, and ∗∗∗ represent p<0.05, p<0.001, and p<10^−5^.

We performed calculations for genetic and age statistics of the model (Theory Supplementary Note). We emphasize that these calculations’ purpose is not precise numerical prediction but rather conceptual investigation of relationships between mutations and age. We observed that: (I) The number of cancer-causing mutations per cell in normal tissue grows with age (Extended Data Figure 6A). (II) Cancer incidence rises with age (Extended Data Figure 6B). (III) Cancers have more cancer-causing mutations than normal tissue (Extended Data Figure 6A). These first three observations are also predicted by the classical mathematical model of multi-hit carcinogenesis which represents that mutations arrive randomly in a tissue and that the *k*th mutation along a lineage defines the initiation of cancer^70,71^. The key distinction between our formulation and the classical model is that we allow that mutations may vary in their carcinogenic strength. Our model therefore makes three further predictions that the classical model does not. (IV) The number of cancer-causing mutations per cancer grows with the age of cancer initiation (Extended Data Figures 6A,D) (V) Cancers with fewer cancer-causing mutations tend to have mutations with stronger carcinogenic effects (Figure 6B). (VI) Cancers at younger ages tend to have mutations with stronger carcinogenic effects than cancers at older ages (Figures 6C, Extended Data Figure 6E).

Predictions (I) to (III) are consistent with extensive data across cancer and tissue types^12,16,36,51,71–74^. Prediction (IV) is consistent with the positive correlation between patient age and number of non-synonymous mutations among putative driver genes that we observed for many cancer types, although uterine corpus endometrial carcinoma is a notable exception (Extended Data Figure 6F). Prediction (V) is consistent with AML, COADREAD, and ESSC data (the three cancer types whose available normal tissue data enabled estimation of carcinogenic effects): the mean carcinogenic effect of driver mutations per patient negatively correlates with patient age (Figure 6D). Prediction (VI) too is consistent with AML, COADREAD, and ESSC data: the mean carcinogenic effect of drivers per sample negatively correlates with the number of driver mutations (Figure 6D). Thus, a simple set of postulates – namely, mutations arrive stochastically and vary in their carcinogenic strength – is already enough to qualitatively predict multiple observations in cancer and normal tissue data. Overall, these results support that the most powerful cancer-causing mutations cut down a cell’s mutational and temporal distance to cancerous transformation, and so these mutations may be found preferentially in the simplest cancer genomes and at the youngest patient ages.

## DISCUSSION

In the search for cancer-causing mutations, tens of thousands of cancer genomes have been sequenced, and signals of positive selection have been identified. These signals’ interpretation needs revisiting, however, in the light of modern findings that positive selection is ubiquitous in our healthy tissues. Cancer causation needs to be conceptually and inferentially distinguished from positive selection.

Towards one possible solution, we have defined a mutation’s carcinogenic effect in terms of the per cell rate of cancerous transformation. Our metric is simplistic. The multi-stage evolutionary process of carcinogenesis and its combinatorial genetic interactions are boiled down, by our metric, to a single number per alteration. This one-dimensional probabilistic assessment of alterations’ cancer-causing power has the disadvantage of missing innumerable biological details; on the other hand, its minimalism brings the benefits of clear interpretation and broad application across cancer types and mutations.

What’s the purpose of assessing mutations’ carcinogenicity? Beyond the basic question of cancer’s origins, there may be implications for the modulation of cancer risk. For a mutation with known carcinogenic effect, perturbation of the mutation’s frequency has a predictable consequence on a tissue’s risk of cancer (Figure 1B). Perhaps someday soon, preventative treatments will reduce the frequencies or inhibit the effects of carcinogenic mutations in our tissues before cancer develops. Conversely, it is thought that exogenous carcinogens increase cancer risk largely by elevating frequencies of carcinogenic mutations in our tissues, not only through mutagenesis but also by exerting selective pressures, for example, chemotherapies cause expansion of *TP53* mutations in the blood^38^, while air pollution is associated with expansion of *EGFR* mutations in the lungs^75^. Broadly therefore, for mutations across tissues, estimating carcinogenic effects could contribute to our understanding and management of cancer risk in the population.

Carcinogenic effect estimates may also help to establish priority mutational targets for cancer treatments. Mutations in cancers, regardless of their carcinogenic effects, can in principle be therapeutically targeted to identify and kill cancer cells. Yet, treatment resistance is a pervasive problem. We hypothesize that treatment resistance can be reduced by targeting highly carcinogenic mutations; conceivably, cancers cannot easily evolve absence of such mutations’ pathways without losing the cancerous phenotype. This hypothesis should be testable in the future by comparing clinical data against targeted mutations’ carcinogenic effect estimates.

We have estimated carcinogenic effects for mutations in the blood, colon, and esophagus. Underlying our inferences, the basic idea is that somatic evolution generates genetically diverse tissues, providing a testing ground for mutations’ carcinogenicity. This natural experiment’s major limitation, however, is its visibility at epidemiological scale: for most normal tissues, sequencing studies have small cohort sizes, restricting the resolution with which population-wide mutation frequencies are estimated. We therefore sought to infer carcinogenicity from cancer genomes without the normal tissue reference.

We have proposed that patient age data can contribute key, previously underappreciated, causal information to cancer genomic analyses. Patient ages alongside cancer genomes can test the null hypothesis that mutations are not carcinogenic. Age data moreover provide information on the magnitudes of mutations’ cancer-causing strength. Indeed, it stands to reason that age data can aid interpretations of cancer genomes in terms of the somatic evolutionary ageing process that generated the genomes. Yet, age has not been regarded sufficiently important to be included in many cancer genetic datasets; ∼30% of patients in COSMIC are missing age annotations.

Interestingly, the degree to which age distributions differentiate purported driver genes varies widely between cancer types (Figure 3A). AML and glioblastoma show the greatest age differentiation while epithelial cancers show the least. One explanatory factor underlying these cancer type-specific age distributions appears to be magnitudes of mutations’ cancer-causing strength: our carcinogenic effect estimates are at least an order of magnitude greater for AML than esophageal and colorectal cancers (Figures 1D,1E, Extended Data Figure 1E). A second possible contributing factor could be that for some cancer types more than others, mutations’ carcinogenic effects depend on age-specific tissue biology, though for AML, perhaps surprisingly, we have estimated that mutations’ carcinogenic effects differ between younger and older patients far less than mutation frequencies differ between the age groups (Figures 3C,D), suggesting that AML etiology may be simpler than previously thought. A third factor potentially contributing to cancer type-specific age distributions could be tissue structure. In the blood, clonal expansions follow exponential growth curves^15,40^; for solid tissues on the other hand, spatial structure may restrict clonal expansions to sub-exponential growth curves^57^, which according to theory diminishes age divergence between driver mutations (Extended Data Figures 4A,B). Fourthly, for some tissues, mutation frequencies are substantially shaped by environmental insults, weakening the statistical relationship between genetics and chronological age^76^. External exposures could thus help to explain why age dependence of driver mutations is less for epithelial cancers than for AML and glioblastoma, the latter two cancers arising from stem cell lineages comparatively protected from the outside world. For the future, the lifelong somatic evolutionary trajectory from zygote to cancer, and its mutational and environmental causes, will be increasingly clarified by the growing documentation of the aging genetic landscapes of our normal and pre-cancerous tissues, ultimately informing cancer prevention.

## METHODS

### Data sets

For each cancer type that we analyzed, we obtained some or all the mutation data from the pan cancer whole exome sequencing database that is COSMIC version 98^41^. This database is a collation of many cancer genomic studies and the classification of some cancer types requires clarification. A curated subset of the COSMIC dataset is the TCGA dataset which has a standardized classification scheme. We used the TCGA classification of cancer types, which we extended to the whole COSMIC dataset as follows: for each cancer sample in COSMIC but not TCGA, if the sample’s anatomical site and histology matched that of samples in TCGA with a specific cancer type classification, then we assigned that classification to the sample, otherwise we disregarded the sample. For the cancer types in our cancer type-specific analyses, namely AML, GBM, ESSC, COADREAD, and BRCA, we performed a manual review of the studies that contributed to the COSMIC database to ensure correct classifications. We disregarded a small subset of studies where the cancer type classification was ambiguous or erroneous. We also disregarded samples with no patient age annotation. For our pan cancer COSMIC analyses (Figure 3A, Extended Data Figure 6D), we focused on cancer types with at least 300 samples. Our filtered version of the COSMIC data is available at https://github.com/dmcheek/cancer-mutations.

For mutations in normal blood, we used two datasets. The first is Supplementary Table 2 of Abelson et al.^39^, comprising 817 normal blood samples from 676 donors who did not develop any hematological malignancies within ∼10 years of sampling. 141 of the donors had multiple blood samples available, spanning a median of 10.5 years. The donors are aged between 36 and 90 years. Targeted deep sequencing of AML-associated genes was performed, with minimum reported VAF 0.001. The second dataset is Supplementary Table 2 in Fabre et al.^40^, comprising 1,593 blood DNA samples from 385 donors aged between 54 and 103 years at the time of sampling. The participants had no history of hematological malignancy. Their blood was sampled up to 5 times each, over a period of 3–16 years. Targeted deep sequencing of AML- and clonal hematopoiesis-associated genes was performed with mean coverage ∼1000x and minimum reported VAF 0.0002.

For AML mutations, we appealed to four datasets. One, Supplementary Table 3 of Tazi et al.^43^ presents sequencing data for 2113 AML patients enrolled in UK-NCRI trials^77–79^, aged between 15 and 100 years. Two, we obtained data from http://www.cbioportal.org/study/summary?id=aml_target_2018_pub on somatic mutations of 886 AML patients between 0 and 30 years old, the data being part of a pediatric AML trial of the Children’s Oncology Group, National Cancer Institute^55^. Third, we obtained data from http://www.cbioportal.org/study/summary?id=aml_ohsu_2022 on 742 AML patients aged from 0 to 80 years, as part of the Oregon Health & Science University Beat AML cohort^42^. Fourth, we obtained AML mutation data from COSMIC, comprising 908 AML patients ranging from 0 to 88 years old. Together, these 4 AML and 2 normal blood datasets include a variety of ranges of the genome sequenced, including the whole genome, the whole exome, and specific genes. We focused on 21 genes that were covered by all the AML and normal blood studies and were listed in Bailey et al.^44^ as AML driver genes.

For our comparison of driver gene mutations between esophageal squamous cell carcinoma and normal esophageal epithelium, we used data from Yokoyama et al.^17^, presented in their Supplementary Table 8. The cohort comprises 68 ESSC patients. The patients are aged between 47 and 79 years. Each patient donated one ESSC sample and one ‘physiologically normal’ epithelium sample. The biopsy area is ∼8mm^2^. See their Supplementary Table 2 for donor and sample information. 24 genes were sequenced. The sequencing depth was not specified though the minimum VAF reported is 0.05. For a fair comparison of cancer and normal tissue, we wanted to avoid the scenario that the normal tissue samples are part of a clone that gave rise to the cancer, and so we removed 5 patients who had mutations in common between their cancer and normal tissue samples. We then focused on the subset of genes with signals of positive selection in ESSCs, as declared in Supplementary Table 5 of Yokoyama et al.^17^.

For mutations in the normal colon, we turned to Lee-Six et al.^51^ and Olafsson et al.^14^, the former including 42 individuals without diagnosed inflammatory bowel disease and the latter comprising 28 ulcerative colitis patients and 18 Crohn’s disease patients, with ages ranging from 11 to 80 years. Both studies performed whole genome sequencing on single colonic crypts, and coding mutations were collated in Supplementary Table 3 of Olafsson et al.^14^, excluding one patient from Lee-Six et al.^51^ whose colon was heavily mutagenized by prior chemotherapy. In total, the data comprise 1395 crypts among 91 donors.

We obtained data on cytogenetic abnormalities among 2113 AML samples from Tazi et al.’s^43^ Supplementary Table 4, representing the same patients whose driver gene mutations we analyzed. The chromosomal variants declared include amplifications and deletions of whole chromosomes and chromosome arms, as well as inversions and translocations. Copy number neutral loss of heterozygosity variants are not included in this AML dataset. We obtained data on amplifications and deletions at the level of whole chromosomes and chromosome arms among 482,789 normal blood samples in the UK Biobank from https://github.com/blundelllab/mCA-mutation-rates-fitness-consequences ^54^; these copy number alterations were originally called by Loh et al.^67^, and they include estimates of the fraction of blood cells per sample with each alteration. The donor ages range from 40 to 75.

We obtained copy number annotations for the TCGA pan-cancer dataset from https://github.com/VanLoo-lab/ascat, and here we focused exclusively on those samples which the authors specified to be without whole genome doubling. Then to map their copy number annotations to the list of putative oncogenes and tumour suppressor genes from Bailey et al.^44^, we used genomic coordinates for genes from https://useast.ensembl.org/index.html. No declared copy number alteration has a genomic boundary that lands within a putative driver gene; that is, each copy number alteration unambiguously covers a specific gene or does not cover that gene.

### Estimating mutant cell fractions in bulk normal tissue

We focused on non-synonymous SNVs and small indels. For each observed mutation in a normal esophagus or blood sample, we multiplied the variant allele frequency by two to estimate the fraction of cells in the sample with the mutation. This estimate is based on the simplifying assumption of diploidy and heterozygosity. We adjusted the estimates in two special scenarios: Firstly, for some mutations in some samples, the VAF multiplied by two exceeded one, which we interpreted as possible loss of heterozygosity, and so we set the mutant cell fraction estimate equal to the VAF without multiplying by two. Secondly, for males and the X chromosome, we again set each mutation’s cell fraction estimate equal to the VAF. Then we extended our mutant cell fraction estimates to the level of genes. For this, calculation elaboration is needed where there are multiple mutations per gene per sample. To estimate the fraction of cells in a sample with a specific gene mutated, we took the *summation* of the gene’s mutations’ cell fraction estimates, unless this summation exceeded one, in which case we took the *maximum* of the mutations’ cell fraction estimates. The former case represents the assumption that the gene’s mutations land in distinct cells within a sample, while the latter case represents the assumption that the mutations land in the same cells. These estimates should be regarded as ballpark figures, since mutational cooccurrences per cell, copy number status, and sample purity are unknown, and mutations at low VAF (<0.05 for the esophagus and <0.001 for blood) are unreported. Next, for each gene, we took the mean across samples of the per sample mutant cell fraction estimates, thereby obtaining estimates for the fraction of cells mutated per gene in the cohort. Here, to assess sampling noise (as opposed to uncertainty introduced by measurement or processing errors), we calculated 95% bootstrap confidence intervals for these cell fraction estimates: for each of 1000 bootstrap resamples of the tissue samples, we calculated mean mutant cell fractions per gene, thereby generating a gene-specific bootstrap distribution for the mean mutant cell fraction, and the 2.5 and 97.5 percentiles of the distribution define the confidence interval. We note the theoretical caveat that bootstrap intervals in general struggle to quantify sampling noise for small cohort sizes.

### Carcinogenic effect estimates

The ESSC carcinogenic effect estimates are based on the patient-matched ESSC and normal esophageal epithelium data from Yokoyama et al.^17^. The adult AML carcinogenic effect estimates are based on the normal blood data from Abelson et al.^39^ and Fabre et al.^40^ and the AML data in COSMIC, Bottomly et al.^42^, and Tazi et al.^43^, with a focus on AML patients older than 36 years old (since the youngest normal blood donor is 36). For each driver gene, we compared the fraction y of cancer samples with the gene mutated (non-synonymous SNVs and indels) against the estimated fraction x of normal tissue cells across the cohort with that gene mutated: the odds ratio 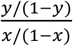 is the carcinogenic effect estimate. See the Theory Supplementary Note for a critical discussion of the assumptions that underly this estimate. We performed 1000 bootstrap resamples of the donors in each of the cohorts to obtain 95% bootstrap confidence intervals for the carcinogenic effect estimates.

For driver gene mutations in the older AML group, we also provided an alternative approach to estimate carcinogenic effects for the older AML group, with the purpose to more clearly account for age in the comparison of AML and normal blood mutations. For this, we employed a hybrid regression model with four real-valued gene-specific parameters *a, b, ϕ, w*. The model’s first component is that for an AML sample at patient age *t*, a gene is mutated with probability 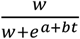. Here, *w*> 0 is the carcinogenic effect and 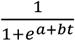 is the expected fraction of normal blood cells with the gene mutated. The model’s second component is that for a normal blood sample at age *t*, the fraction of cells with the gene mutated follows a Beta distribution with shape parameters 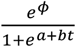 and 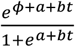; so the mean of the distribution is 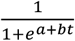 while the spread of the distribution is controlled by *ϕ*. To address the invisibility of mutations at low cell fractions, we then specified that the observed mutant cell fraction in a normal blood sample is distributed as *X* × *I*(*X* ≥ *x*_0_), where 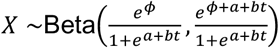 is the actual mutant cell fraction, *I* is the indicator function, and *x*_0_ denotes the minimum cell fraction estimate among observed mutations. As such, we obtained a likelihood function for mutation frequencies in the AML and normal blood data, with which we performed maximum likelihood estimation for each gene independently. The carcinogenic effect estimates from this regression model strongly agree with the carcinogenic effect estimates from the unadulterated odds ratio of mutation frequencies between adult AML and normal blood (rho=0.98, p=4.8e-06).

To estimate carcinogenic effects for the younger AML group (<25 years) in the absence of age-matched normal blood data, we used the same data as for the adult AML analysis, supplemented with the childhood AML data from Bolouri et al.^55^. We began with the simplifying approximation that a normal HSC carries a mutation within a specific gene with probability proportional to the gene’s length: that is, for a gene with length *L*, the gene is on average mutated in fraction *cL* of a child’s HSCs, for some unknown constant *c*. Then, supposing that a mutation in this gene confers a *w*-fold elevation in the rate of cancerous transformation, the gene is expected to be mutated in fraction *f* = *wcL*/(1 − *cL* + *wcL*) of AMLs. Approximating further that *cL* is small relative to one, *wc* = *L*^−1^*f*/(1 − *f*). By the functional invariance of maximum likelihood estimation, the maximum likelihood estimate for the relative carcinogenic effect *wc* is given by 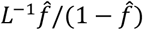, where 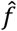 is the observed fraction of samples mutated. We obtained 95% binomial confidence intervals for *f*, which we transformed via the equation *wc* = *L*^−1^*f*/(1 − *f*) to obtain 95% confidence intervals for the relative carcinogenic effect. We note an important methodological distinction between the childhood and adult AML carcinogenic effect estimation: for childhood AML, the normal blood benchmark is formed by gene lengths who approximately represent the total number of possible mutations in a gene, whereas for adult AML, the normal blood benchmark is formed by bulk DNA samples who exhibit only those mutations at variant allele frequency high enough to be distinguished from sequencing errors, so the adult carcinogenic effect estimates refer primarily to those mutations in a gene who drive expansion in normal blood whereas the childhood carcinogenic effect estimates refer to the aggregate of all mutations per gene. More broadly, we expect the age group-specific modeling assumptions to drive divergence between childhood and adult AML carcinogenic effect estimates, suggesting that Figure 3D understates the real biological agreement between the age groups.

To estimate carcinogenic effects for colorectal cancer, we used cancer data from COSMIC and normal colon data from Lee-Six et al.^51^ and Olaffson et al.^14^ We employed a logistic regression model: for each colorectal cancer driver gene listed in Bailey et al.^44^, we modeled that the gene’s likelihood of being mutated in a sample takes the form 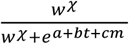, where *t* is the individual’s age, *m* is the exonic mutation burden of the sample, *χ* takes the value 1 or 0 to respectively represent that the sample is a colorectal cancer or a normal colonic crypt, and *a, b, c, w* are the model’s parameters. We calculated maximum likelihood estimates for the carcinogenic effect *w* along with 95% profile confidence intervals.

### Statistics for the age distribution of cancer mutations

For each cancer type and driver gene, we calculated the mean age of patients in the cohort weighted by the number of non-synonymous substitutions and small indels in the gene, along with 95% Wald confidence intervals for this mean age statistic. To assess the degree of age divergence between driver genes, we then calculated the variance between driver genes of the mean age statistic, weighted by the number of mutations per gene, which we divided by the variance of the patient age distribution per cancer type. To assess whether the age divergence between driver genes is significantly greater than zero for each cancer type, we calculated a p value from a Kruskal-Wallis test of the null hypothesis that mutations among driver genes have patient ages that are independent of the identity of the driver genes in which they reside. We also calculated the mean age of non-synonymous substitutions and small indels per driver gene minus the mean age of exome wide synonymous mutations for each cancer type; to generate a null distribution, we recalculated this statistic for each of 1000 permutations of patient ages among samples of the cancer type in question. For these comparisons of driver gene mutations’ age distributions against synonymous mutations, we focused on samples in COSMIC whose mutation burden is no greater than 10 times the median mutation burden per cancer type, to avoid that statistical signal is diminished by a small number of samples with extreme mutation burden.

To compare the age distributions of SCNAs vs. SNVs/small indels, we introduced the *age bias* statistic, defined by the rank biserial correlation between the ages of patients with vs. without a particular alteration. Specifically, for each driver gene and cancer type, among cancer samples with no SNCA in that gene, we calculated the rank biserial correlation between the ages of patients with vs. without a non-synonymous SNV/small indel in the gene; separately, among cancer samples without a non-synonymous SNV/small indel in that gene, we calculated the rank biserial correlation between the ages of patients with an amplification of the gene vs. no SCNA in that gene, and we calculated the same statistic for deletions. We used the r package “effectsize” for point and interval estimates of the rank biserial correlation (Supplementary Table 2).

### A branching process model for a mutation’s trajectory in somatic evolution

Our simplest model of somatic evolution, exhibited in Figure 2, illustrates the stochastic acquisition of some mutation z in (a patch of) normal tissue, where the mutation may alter cells’ rates of division, death, and carcinogenesis. We assume that cells without mutation z divide and die at an equal rate, so for simplicity, we approximate their total number in normal tissue to be some constant *N*_0_. Then the number of cells with mutation z follows a continuous-time Markov process with transitions *i* → *i* + 1 at rate *N*_0_*v* + *ia*_1_ and *i* → *i* − 1 at rate *ib*_1_. Here, *v* is the per cell rate of mutation z acquisition, *a*_1_ is the division rate of cells with mutation z, and *b*_1_ is the death rate of mutant cells. Mutation z’s selective effect in normal tissue is *s* = *a*_1_ − *b*_1_. At time *t*, the rate of cancer initiation per cell is *r*_1_(*t*) given mutation z and *r*_0_(*t*) in the absence of mutation z. Mutation z’s carcinogenic effect is the ratio *r*_1_(*t*)/*r*_0_(*t*). We now explain how to calculate the overall rate of mutant-z cancer arrival, which requires further notation. Let *p*_*k*_(*t, T*) denote the probability of no mutant-z cancer between times *t* and *T* conditional on *N*_1_(*t*) = *k*. Observe the backwards Kolmogorov equations

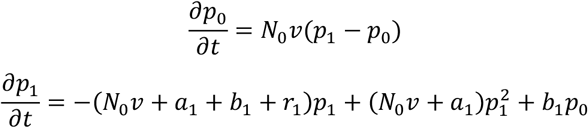

with the boundary condition *p*_*k*_(*T, T*) = 1. We differentiate the backwards Kolmogorov equations with respect to *T* to obtain

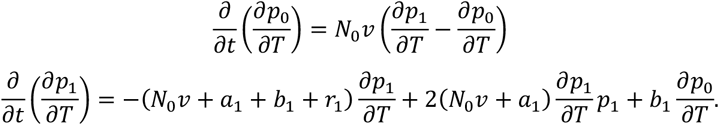

Also note the boundary condition 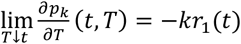. The differential equations can be solved numerically via the Euler method. Then, conditional on no cells with mutation z at time 0 and no cancer between 0 and *T*, the rate of mutant-z cancer at time *T* is 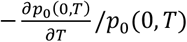 whereas the rate of wildtype-z cancer at time *T* is 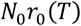. Therefore, for cancer arising at time *T*, the odds that it carries mutation z is 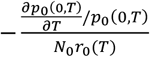. To express the same quantity in more probabilistic notation, let *E* denote the expectation operator, then 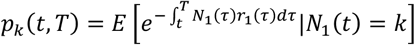 and 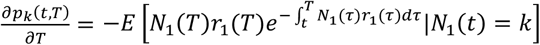, so cancer at time *T* carries mutation z with probability 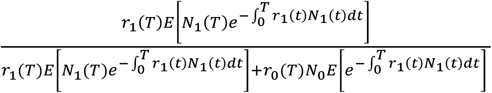 and odds 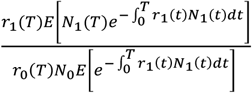. Meanwhile, given absence of cancer by time *T*, the odds for mutation z in a normal tissue cell is 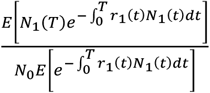 Observe that the odds of the mutation in cancer divided by the odds of the mutation in a normal tissue cell at a given age is precisely the mutation’s carcinogenic effect.

### Null hypothesis of no carcinogenicity

While classically, a cancer driver gene is identified by elevated frequencies of functional relative to neutral mutations, we proposed to additionally compare the age distribution of functional mutations against that of neutral mutations: we proposed that elevated frequencies AND younger ages of non-synonymous relative to synonymous mutations in cancer genomes constitutes evidence for carcinogenicity. This test is based on minimal modelling assumptions: mutations arrive as a Poisson process in normal tissue according to the infinite sites assumption, a neutral mutation on average does not change in variant allele frequency, a positively selected mutation on average increases in variant allele frequency, and a non-carcinogenic mutation’s probability of being in a cancer founder cell is equal to its frequency in the background normal tissue. In the Theory Supplementary Note, we express these assumptions in mathematical notation, and we prove that positively selected, non-carcinogenic mutations have an age distribution that stochastically dominates the age distribution of neutral mutations.

### Multi-hit model of carcinogenesis

Our model of multi-hit carcinogenesis in Figure 6 disregards selection in normal tissue. It specifies that mutations arrive as a rate *v* Poisson process along a normal tissue lineage according to the infinite sites assumption, and that each mutation has a carcinogenic effect independently sampled from a Gamma distribution. So, a lineage at age t has a Poisson distributed number of mutations with mean vt, and the product of these mutations’ carcinogenic effects multiplied by some baseline rate is the rate of carcinogenesis. Calculations for the joint distribution of mutations’ carcinogenic effects and patient ages as well as the number of mutations per cancer are presented in the Theory Supplementary Note, where other models of somatic evolution towards carcinogenesis are explored too.

## Supporting information

Theory Supplementary Note

## Data and code availability

Code generated during this study is available at https://github.com/dmcheek/cancer-mutations.

## Supplemental information

Theory Supplementary Note (pdf) Supplementary tables (excel):

- Supplementary Table 1: Statistics for driver gene mutations in COSMIC
- Supplementary Table 2: Statistics for SCNAs and driver gene mutations in TCGA
- Supplementary Table 3: Cancer type acronyms

## Acknowledgements

This work was supported by the National Institutes of Health (R37CA225655, R01CA279054, R01CA26928 and P01HL142494), by the Glenn Foundation for Medical Research and by an Emerging Leader Award from the Mark Foundation for Cancer Research.

## Author contributions

D.C. developed the conceptual framework and performed analyses. M.B. helped to process data. M.B., M.N., and T.A. provided key advice. K.N. supervised the project. D.C. and K.N. wrote the manuscript.

---

**Extended Data Figure 1:**
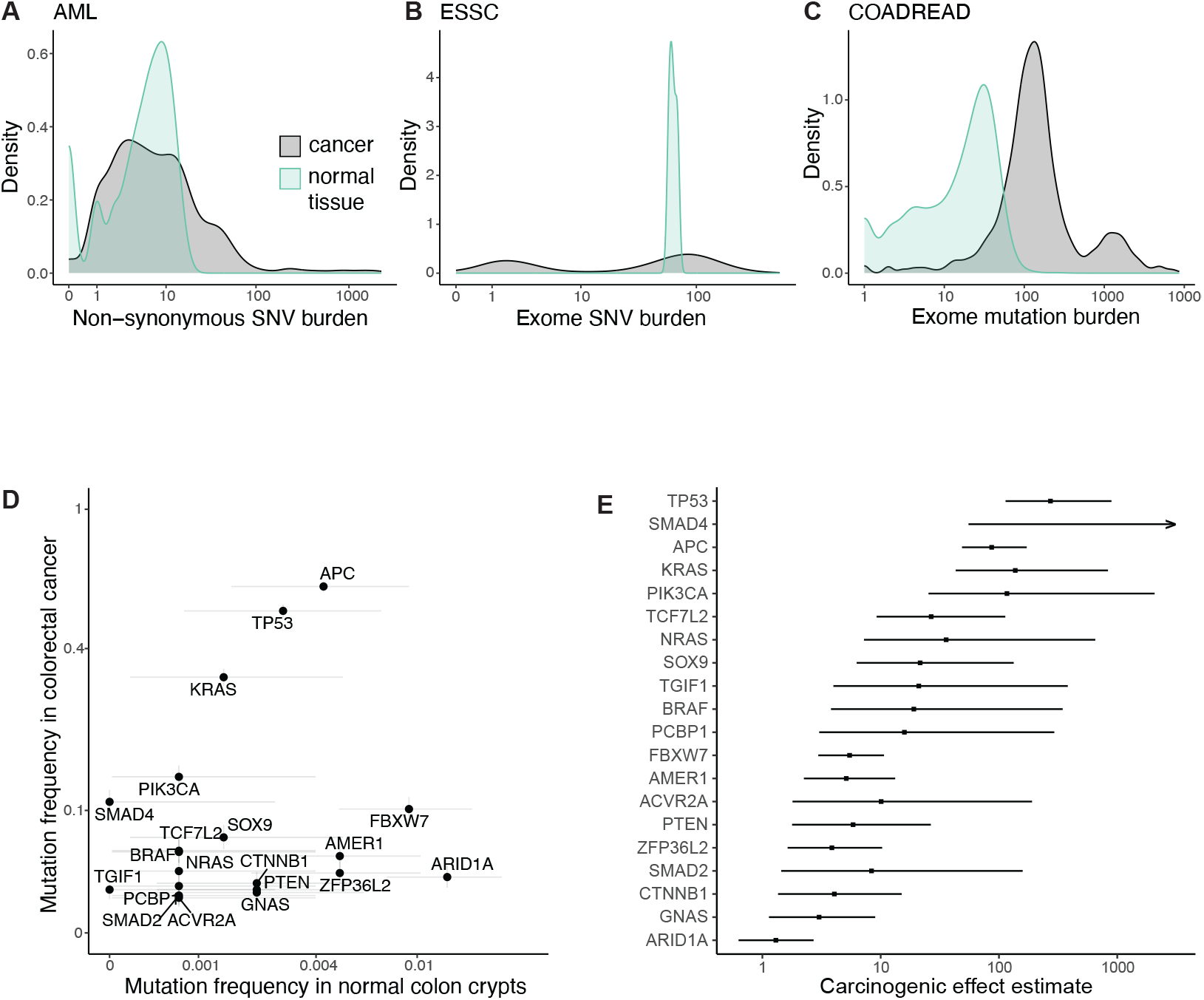
Mutation burden comparisons and carcinogenic effect estimates for colorectal cancer. (A) Non-synonymous SNV burdens in 1224 AML samples from COSMIC and 3592 single normal hematopoietic cell-derived colonies from Mitchell et al.^50^. (B) Exome-wide SNV burdens in 1509 ESSC samples from COSMIC and 3 single normal esophageal epithelial cell-derived colonies from drinkers and smokers in Yokoyama et al.^17^. (C) Exome-wide SNV+indel burden distribution in 2411 COADREAD samples from COSMIC and 1387 normal colonic crypts, from donors with and without inflammatory bowel conditions, in Lee-Six et al.^51^ and Olafsson et al.^14^. Mutational types differ between (A), (B), and (C) due to data accessibility. (D) Mutation frequencies among COADREAD driver genes (Bailey et al.) in COADREAD samples and normal crypts. Samples as in (C) but excluding 787 COADREAD samples whose exonic mutation burden is greater than 251 (maximum of normal crypt mutation burdens). Error bars are 95% binomial confidence intervals. (E) Carcinogenic effect estimates for COADREAD driver genes with 95% Wilks confidence intervals, based on logistic regression model that accounts for mutation burden and patient age (Methods).

**Extended Data Figure 2:**
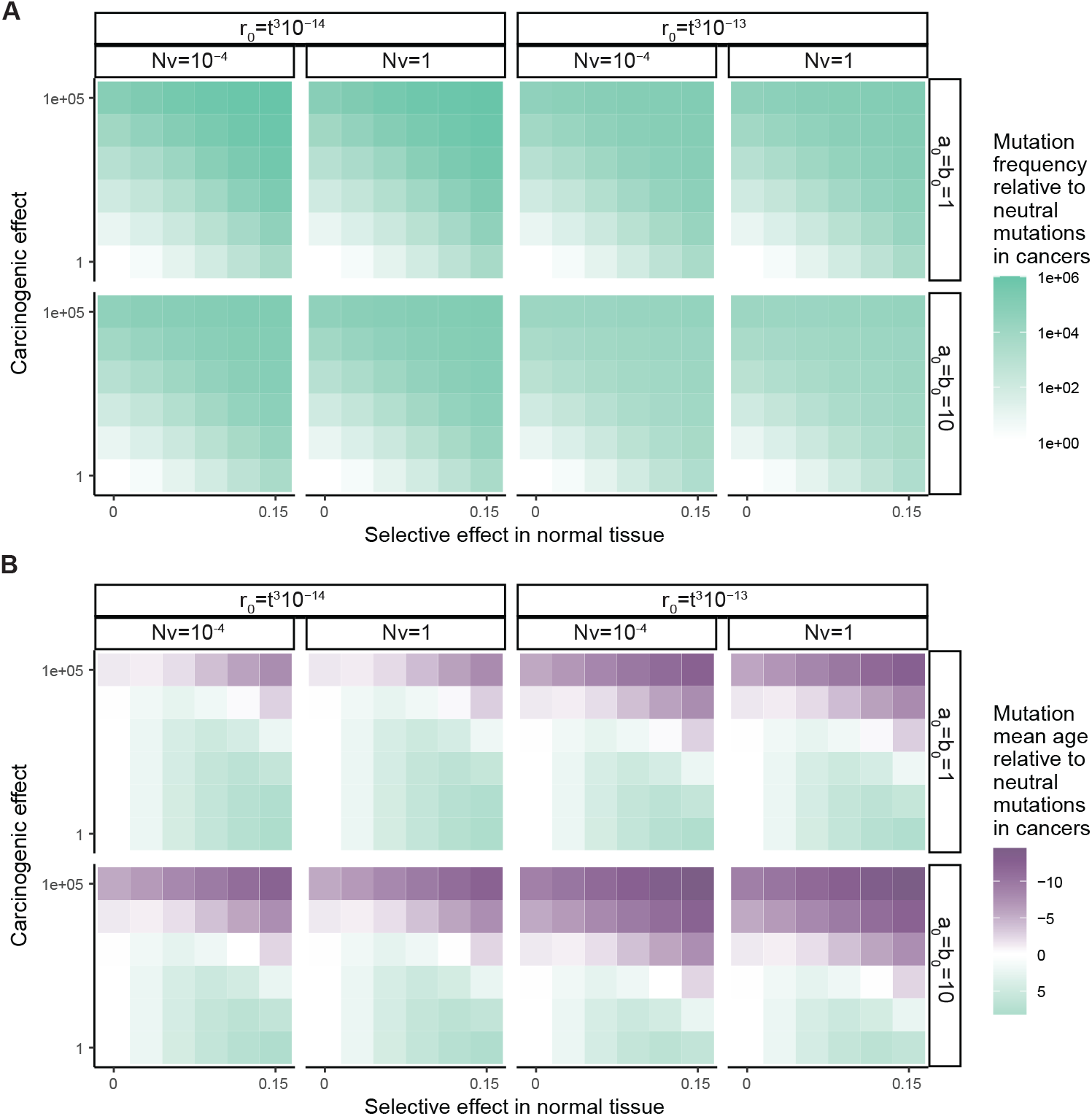
Parameter space exploration for branching process calculations. (A,B) Calculations for model of Figure 2. (A) Frequency of cancers with mutation z divided by frequency of cancers with neutral mutation of same acquisition rate. (B) Mean patient age for cancers with mutation z minus mean patient age for cancers with neutral mutation of same acquisition rate. (A,B) Model and parameter notation as in Figure 2.

**Extended Data Figure 3:**
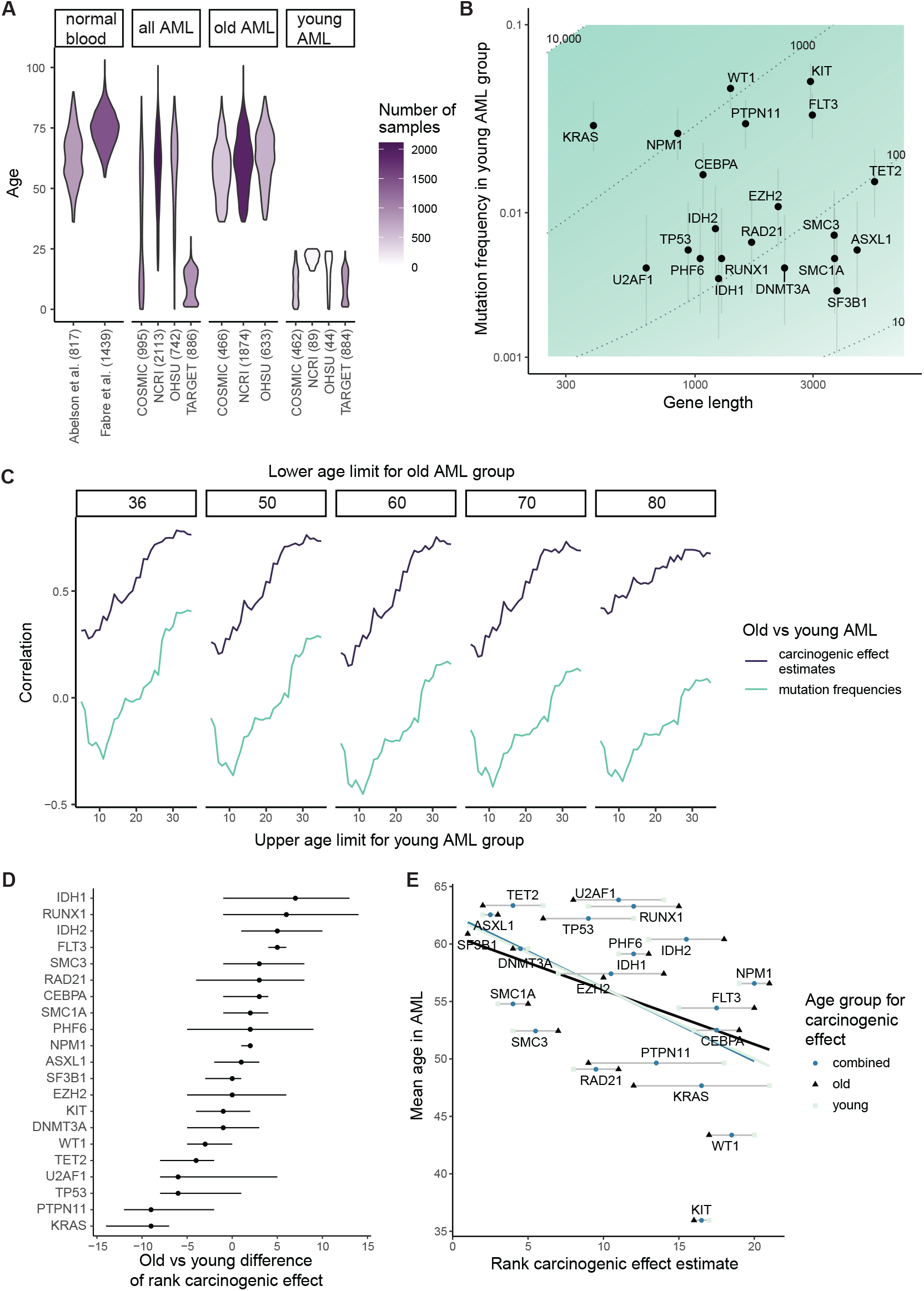
Supplementary analyses on AML. (A) AML data overview. Numbers of samples and ages of individuals among normal blood and AML datasets used in our analyses. (B) Mutation frequencies per gene among young-onset AML group vs. gene lengths. Contour lines denote estimates of *relative* carcinogenic effects (carcinogenic effects normalized by median carcinogenic effect in older-onset AML, see Methods). Vertical lines are 95% binomial confidence intervals for mutation frequencies. (C) Robustness analysis for comparison of older- and young-onset AML in Figures 3C,D. Spearman’s correlation between age groups in terms of mutation frequencies (light line) and carcinogenic effect estimates (dark line) among AML driver genes. X-axis specifies upper age limit for patient to be included in younger group while subplot title specifies lower age limit required for patient and normal blood donor to be included in older group. (D) Gene-specific comparison of carcinogenic effect estimates between age groups. Genes are ranked by their carcinogenic effect estimates separately in adult and childhood AML, then difference between rankings is calculated. Error bars represent 95% bootstrap intervals for difference based on 1000 bootstrap resamples of individuals. (E) Mean patient age negatively associates with carcinogenic effect estimate. X-axis shows genes ranked by carcinogenic effect estimates separately for older- and younger-AML (triangle, square), and average of older- and younger-AML rank (circle). Mean patient age correlation with carcinogenic effect estimate is rho=-0.44, p=0.048 for older-AML; rho=-0.50, p=0.023 for younger-AML; rho=-0.51, p=0.018 for average rank.

**Extended Data Figure 4:**
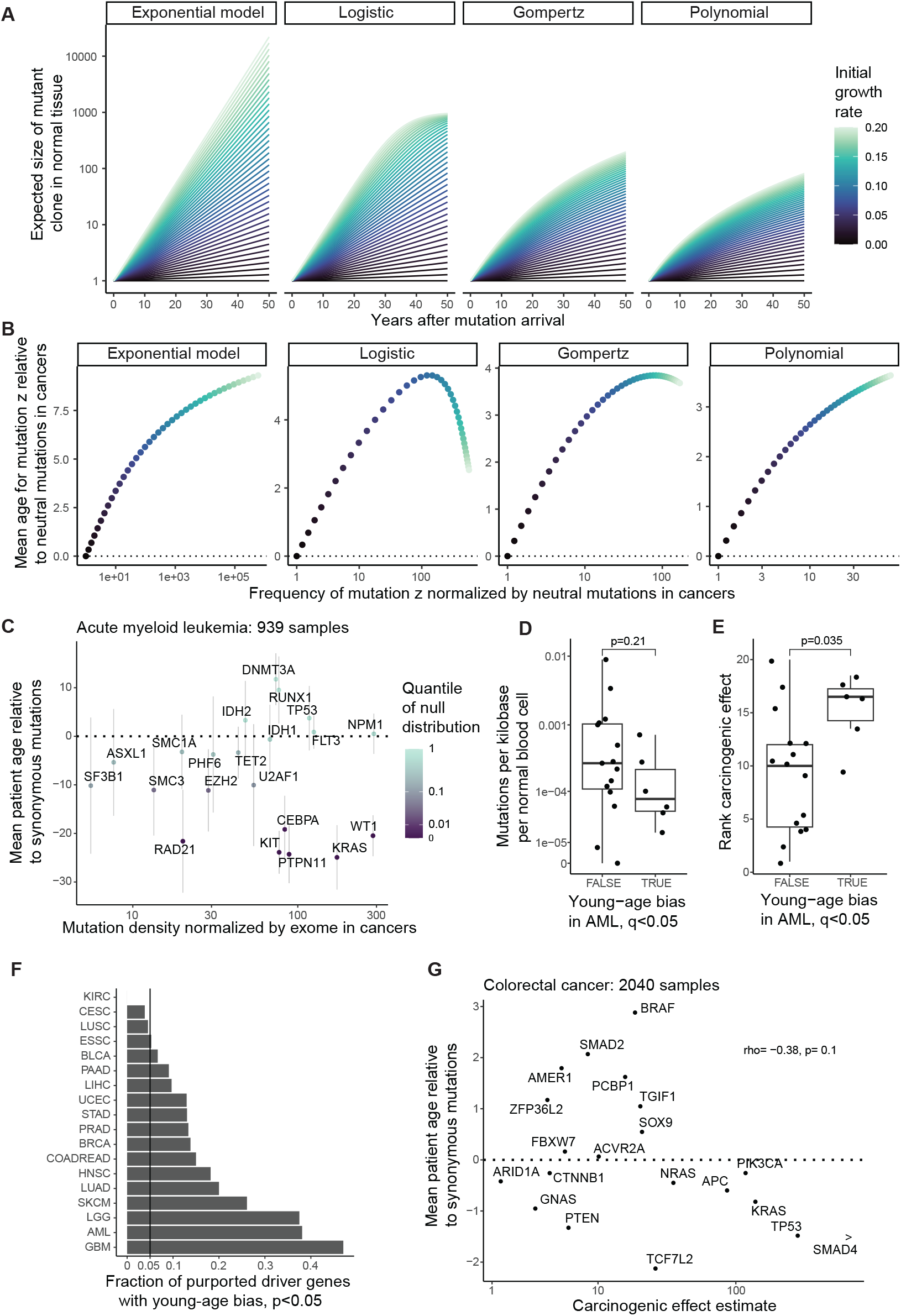
Age distributions of non-synonymous relative to synonymous mutations. (A) Expected growth of positively selected mutant clones in normal tissue, according to exponential, Gompertz, logistic, or polynomial models. (B) Positively selected non-carcinogenic mutations are found preferentially in older patients’ cancers relative to neutral mutations. Y-axis shows the mean patient age for cancers with mutation z minus the mean patient age for neutral mutations. X-axis shows expected frequency of mutation z in cancers divided by expected frequency of neutral mutation with the same acquisition rate. Dot colors correspond to clonal expansion growth curves in normal tissue from (A). (A,B) Full parametrization in Theory Supplementary Note. (C) Null hypothesis test for driver genes in AML. 939 AML samples from COSMIC. Y-axis shows mean patient age for non-synonymous mutations and small indels per driver gene minus the mean patient age for synonymous mutations in exome. Error bars are 95% Wald confidence intervals. Color denotes quantile of age difference statistic with respect to null distribution, generated by permuting patient ages (Methods). X-axis is number of mutations per gene divided by number of mutations in exome across the cohort, normalized by coding sequence length. Genes as in Figures 1E and 3. (D,E) Genes with young-age bias in AML (q<0.05 in C) are less mutated in normal blood (D; Wilcoxon test p=0.21) and have greater carcinogenic effect estimates (E; Wilcoxon test p=0.035). (D) Y-axis is number of mutations in gene per kilobase per normal blood cell, from Figure 3E. (E) Y-axis is rank carcinogenic effect estimate (mean of ranking for young- and older-onset AML). (F) Fraction of driver genes per cancer type with p<0.05, where p is quantile of age difference statistic with respect to null distribution (Methods). Vertical lines denote null expectation 0.05. Cancer types with at least 10 putative driver genes listed in Bailey et al.^44^ and at least 300 samples in COSMIC are plotted. (G) For colorectal cancer, mean patient age relative to synonymous mutations plotted against carcinogenic effect estimates (from Extended Data Figure 1E) per driver gene. *SMAD4* included at finite value for visualization. 2040 colorectal cancer samples in COSMIC.

**Extended Data Figure 5:**
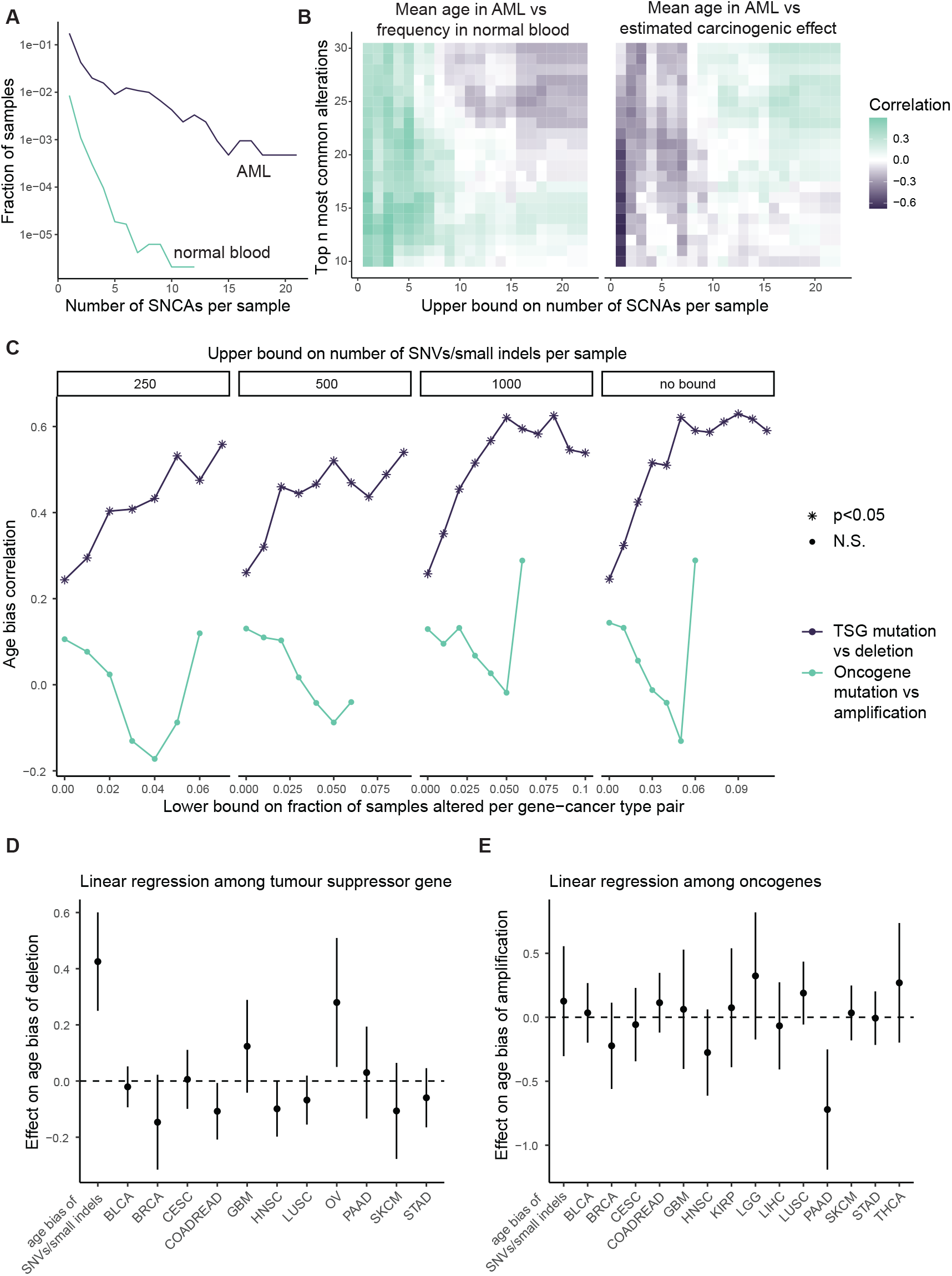
Supplementary analyses for somatic copy number alterations. (A) Fraction of AML and normal blood samples with x autosomal SCNAs, at the level of whole chromosomes or chromosome arms. (B) Robustness analysis for Figures 5B,C. X-axis shows the upper bound filter for the number of SCNAs per sample. Color represents Spearman’s correlation between mean patient age of SCNA vs. mean cell fraction estimate in normal blood (left), or mean patient age vs. carcinogenic effect estimate (right). Correlations are calculated among the top n most frequent SCNAs in the AML and normal blood cohorts, n represented on the y-axis. (C) Robustness analysis for Figure 5D. Subplots show upper bound filter for number of exonic SNVs/small indels per cancer sample. X-axis shows lower bound filter for fraction of samples per cancer type with gene altered; after removing cooccurrences of SCNAs and SNVs/small indels, SCNAs and SNVs/small indels must both exceed lower bound frequency for gene-cancer type pair to be included. Values plotted only if the number of gene-cancer type pairs is at least 20. Y-axis shows Spearman’s correlation among gene-cancer type pairs between age bias of SCNAs vs. non-synonymous SNVs/small indels. (D) Linear regression among gene-cancer type pairs. Age bias of deletions is dependent variable. Age bias of non-synonymous SNV/small indels and cancer type are independent variables. Point estimates and 95% confidence intervals are plotted for the effect of each covariate. (E) Same as (D) but for oncogenes and amplifications. (D,E) Same data as in Figure 5D.

**Extended Data Figure 6:**
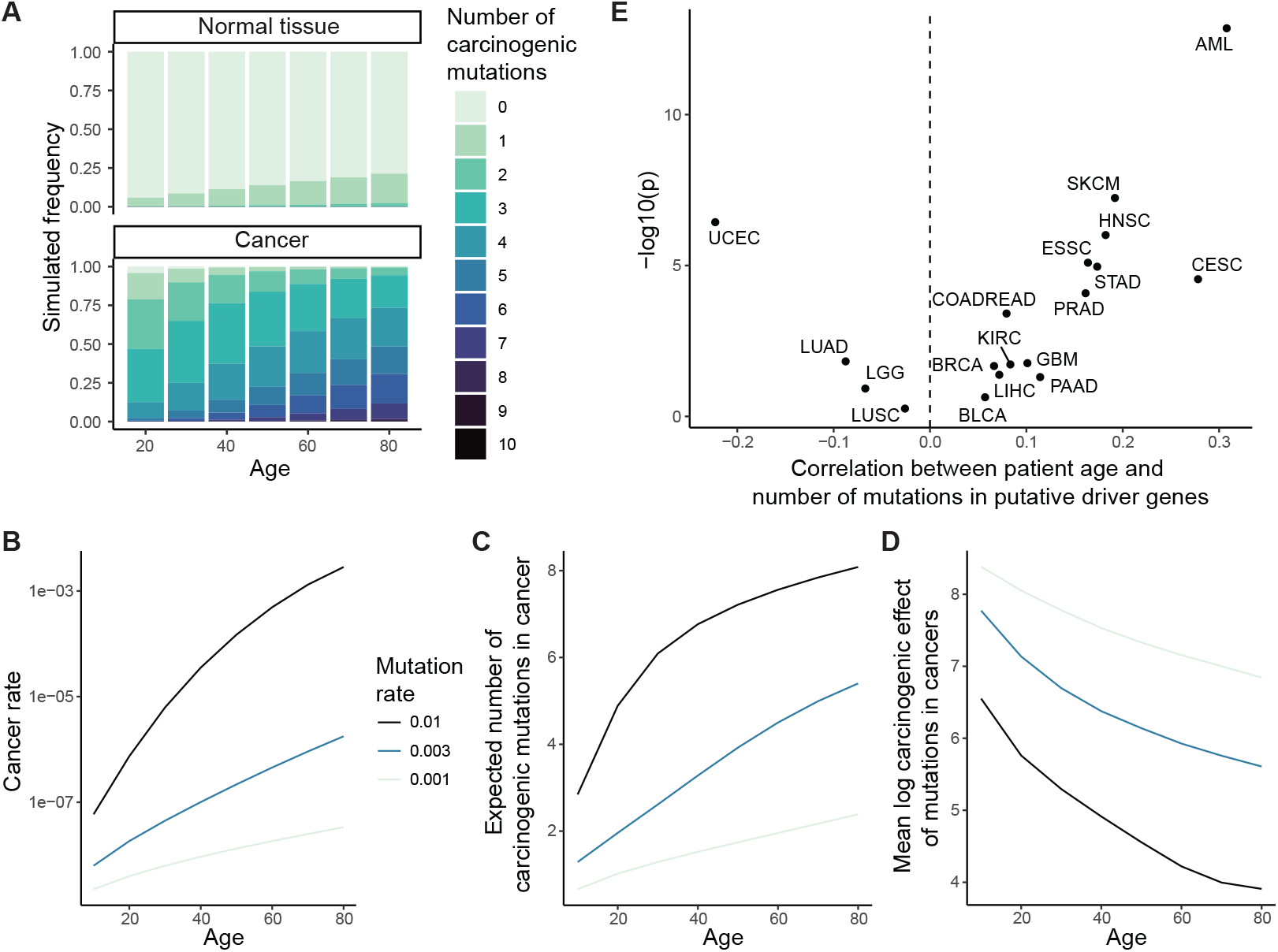
Summary statistics for multi-hit model of carcinogenesis. (A–D) For model defined in Figure 6, (A) numbers of carcinogenic mutations among normal tissue cells and cancers grow with age, (B) rate of cancer initiation per cell per year rises with age, (C) mean number of carcinogenic mutations at cancerous transformation increases with age, (D) mean log carcinogenic effect of mutations at cancerous transformation decreases with age. (A–D) parameters: baseline rate of cancerous transformation without mutations is 10^−9^; mutation acquisition rate along lineage is v=0.003 for (A) or as stated by line color for (B–D); mutations’ carcinogenic effects independently follow Gamma distribution with shape parameter 2 and rate parameter 1. (E) Spearman’s correlation between cancer patient age vs. number of non-synonymous mutations or small indels in putative driver genes. Data from COSMIC. Driver genes listed in Bailey et al.^44^.

