## Supplementary material for "Age distinguishes selection from causation in cancer genomes": Theory Supplementary Note

#### 1 Defining a mutation's carcinogenic effect

In words, we define a mutation's carcinogenic effect as the fold change in the per cell rate of carcinogenesis induced by the mutation. Now we phrase the definition more precisely in mathematical language. Some notation must be established. For a non-cancerous human cell in vivo, we denote the cell's state by an element $(x, y)$ of $\{0,1\} \times \mathcal{V}$, where $x = 1$ represents that the cell has mutation $\xi$ while $x = 0$ represents that the cell does not have mutation $\xi$, meanwhile $\mathcal{V}$ is a space of huge dimension that represents every other factor besides mutation $\xi$ that may be relevant to cancer development, which could for example include other mutations, or the immune system of the person who the cell belongs to. A cell's rate of cancerous transformation is a function $r : \{0,1\} \times \mathcal{V} \mapsto [0,\infty)$ of the cell's state (rate is meant in the sense of a continuous-time Markov process; here there is a simplifying dichotomous assumption that a cell is either cancerous or not). Say that among a cohort of people and within a specific tissue or stem cell lineage/compartment, a uniformly sampled non-cancerous cell has state $(X, Y)$, which is a random variable on state space $\{0,1\} \times \mathcal{V}$. Write $\mathbb{E}$ for the expectation operator. The mutation's carcinogenic effect in the cohort is defined by

$$\frac{\mathbb{E}r(1,Y)}{\mathbb{E}r(0,Y)}. \tag{1}$$

At the end of this document we shall discuss pros and cons of this definition and pose alternative options, but for now let's take (1) as given. Note that the distribution of $Y$ plays a key role in the definition of the mutation's carcinogenic effect: a mutation's effect is context dependent. However, the distribution of $X$ plays no role in the definition: the carcinogenic effect is defined not in terms of the natural state of the mutation but instead by an artificial perturbation of the mutation.

#### 2 Causation, association, and stratification

In contrast to the carcinogenic effect (1), the *observational hazard ratio* for cancer initiation given mutation $\xi$ is defined by

$$H = \frac{\mathbb{E}[r(1,Y)|X=1]}{\mathbb{E}[r(0,Y)|X=0]}, \tag{2}$$

which depends upon the joint distribution of $X$ and $Y$. In other words, $H$ is the ratio of the (per cell) rates of carcinogenesis among cells with vs without the mutation, which is in principle accessible from observational human data. Say that mutation $\xi$ is present in fraction $x$ of normal cells among a cohort of people. Then the mean rates of cancer with and without the mutation are respectively $x\mathbb{E}[r(1,Y)|X=1]$ and $(1-x)\mathbb{E}[r(0,Y)|X=0]$ and, given a large cohort size, the fraction of cancer founder cells that have the mutation is $y = Hx/(Hx+1-x)$. It follows that $\frac{y/(1-y)}{x/(1-x)}$ estimates the observational hazard ratio $H$.

Note that if $X$ and $Y$ are independent, then (2) and (1) are equal. Can (2) therefore act as a proxy for (1)? Well, *association does not imply causation.* This phrase is a religious incantation in science, and deservedly so. There might exist confounding variables. An especially obvious confounding variable here is age. To illustrate, cancer incidence increases with age, as do the frequencies of neutral somatic mutations in our tissues, so a cell's risk of cancerous transformation increases with mutation burden even in the absence of mutational causation of cancer. We seek to ameliorate confoundment by performing stratification. Idealistically, we would like measure some variable $Z$ such that $X$ and $Y$ are conditionally independent given

$Z = z$, in which case the stratified carcinogenic effect

$$\frac{\mathbb{E}[r(1, Y)|Z = z]}{\mathbb{E}[r(0, Y)|Z = z]}$$

would be equal to the stratified observational hazard ratio

$$\frac{\mathbb{E}[r(1, Y)|X = 1, Z = z]}{\mathbb{E}[r(0, Y)|X = 0, Z = z]}. \tag{3}$$

For a down-to-earth impression of the ideal, we compared cancer and normal tissue genomes matched by age and mutation burden to estimate mutations' carcinogenic effects for AML and colorectal cancer. Meanwhile for esophageal cancer, we performed a patient-matched comparison of (lineage-independent) cancer and normal tissue samples. We strongly acknowledge that these comparisons are not immune to confoundment, but we regard them as the best options given available data.

### 3 Age distributions of non-carcinogenic mutations

#### 3.1 Old-age bias for positively selected mutations

For many cancer types, normal tissue study cohort sizes are too limited for precise estimation of (2) let alone (3). We therefore seek assessments of carcinogenicity that do not rely upon normal tissue data. We appeal to cancer genomes and patient age distributions. We propose that an elevated frequency AND a younger patient age distribution for a set of functional mutations relative to neutral mutations in cancer genomes falsifies a null model in which the mutations are not carcinogenic. Our reasoning is simple. Signals of positive selection in normal tissues appear preferentially towards older ages. Therefore signals of positive selection in cancer genomes appearing preferentially towards younger ages cannot be explained by unadulterated inheritance from normal tissue to cancer. To explain such young-age biases we invoke mutations' carcinogenic effects. In this section, we formalise the null model and prove the null prediction that positively selected non-carcinogenic mutations appear in older patients' cancers compared to neutral mutations.

Suppose that there is a set $\{1, 2, .., n\}$ of cancer patients. For patient $i$, $t_i$ denotes their age when cancer initiates in their tissue. In the life history of patient $i$'s tissue, non-carcinogenic mutations arise as a rate $v_i$ Poisson process according to the infinite sites assumption. Each new non-carcinogenic mutation is positively selected with probability $p_+$ or neutral with probability $p_0$. We don't introduce notation for negatively selected nor carcinogenic mutations in this analysis because we don't care about them here. The set of neutral mutations to have arrived in patient $i$'s tissue prior to carcinogenesis is $\mathcal{M}_0^i$ while the set of positively selected non-carcinogenic mutations is $\mathcal{M}_+^i$. Write $\tau_m$ for the (random) arrival time of mutation $m \in \mathcal{M}_+^i \cup \mathcal{M}_0^i$. Write $F_m$ for the fraction of cells with mutation $m$ in the background normal tissue at the time of cancer initiation. We make mild assumptions on the distribution of the $F_m$. For $m \in \mathcal{M}_0^i$, we specify that $\mathbb{E}[F_m|\tau_m = s] = E_0$ does not depend on the cancer initiation time $t_i$ nor the mutation arrival time $s$. This assumption represents the definition of neutrality: a neutral mutation may undergo genetic drift but its *expected* frequency does not change with the progression of time. On the other hand, for $m \in \mathcal{M}_+^i$, we specify that $\mathbb{E}[F_m|\tau_m = s] = E_+(t_i - s)$ is an increasing function of $t_i - s$. This assumption represents the definition of positive selection: a positively selected mutation's expected frequency increases with the time since the mutation's arrival. Also, for simplicity, the $F_m$ are independent (independence could be violated for mutations appearing in the same cell or in the scenario of clonal interference; on the other hand, independence may be reasonable for mutations in distinct regions of a tissue or distinct patients, the latter especially mattering for large cohorts). Let $Z_m = 1$ denote that mutation $m$ is present in the cancer founder and $Z_m = 0$ denote that it is not. The $Z_m$ are independent Bernoulli random variables. The non-carcinogenicity of mutation $m \in \mathcal{M}_0^i \cup \mathcal{M}_+^i$ is represented by the assumption

$$\mathbb{P}[Z_m = 1|F_m = f, \tau_m = s] = f, \tag{4}$$

which says that a mutation's probability of being in the cancer founder cell is equal to its frequency in the background normal tissue. We write $X_+^i = \sum_{m \in \mathcal{M}_+^i} Z_m$ and $X_0^i = \sum_{m \in \mathcal{M}_0^i} Z_m$ respectively for the numbers of positively selected non-carcinogenic mutations and neutral mutations in patient $i$'s cancer. Our main claim is the following.

**Proposition 3.1.** *For $t > 0$,*

$$\mathbb{E}\left[\frac{\sum_{i:t_i>t} X_+^i}{\sum_{i=1}^n X_+^i}\middle|\sum_{i=1}^n X_+^i > 0\right] \geq \mathbb{E}\left[\frac{\sum_{i:t_i>t} X_0^i}{\sum_{i=1}^n X_0^i}\middle|\sum_{i=1}^n X_0^i > 0\right],$$

*i.e. positively selected non-carcinogenic mutations appear in older patients' cancers than neutral mutations.*

Towards proving Proposition 3.1, we use the following elementary result.

**Lemma 3.2.** *Suppose that $\alpha = (\alpha_i : i = 1,..,n)$ and $\beta = (\beta_i : i = 1,..,n)$ are probability distributions with $\alpha_{i+1}/\beta_{i+1} \geq \alpha_i/\beta_i$ for $i = 1,..,n-1$. Then $\alpha$ stochastically dominates $\beta$, that is,*

$$\sum_{i=j}^n \alpha_i \geq \sum_{i=j}^n \beta_i, \quad \text{for } j = 1,..,n.$$

*Proof of Lemma 3.2.* By the monotonicity of $\alpha_i/\beta_i$, there exists $k$ such that $\alpha_i/\beta_i \geq 1 \iff i \geq k$. Then for $j \geq k$,

$$\sum_{i=j}^n \alpha_i \geq \sum_{i=j}^n \beta_i.$$

It remains to prove the result for $j < k$. Since $\alpha$ and $\beta$ are probability distributions, $\sum_{i=1}^n(\alpha_i - \beta_i) = 0$, and therefore

$$\sum_{i=j}^n(\alpha_i - \beta_i) = -\sum_{i=1}^{j-1}(\alpha_i - \beta_i) \geq 0,$$

completing the statement of the lemma. □

Now we return to the main proposition.

*Proof of Proposition 3.1.* From (4), a positively selected non-carcinogenic mutation $m \in \mathcal{M}_+^j$ who arrives in patient $j$'s tissue at time $s$ is bequeathed to patient $j$'s cancer with probability $\mathbb{P}[Z_m = 1|\tau_m = s] = E_+(t_j - s)$. Appealing to the thinning property of Poisson processes, the number of positively selected non-carcinogenic mutations in patient $j$'s cancer, $X_+^j$, is Poisson distributed with mean $v_j p_+ \int_0^{t_j} E_+(s)ds$. Then, $X_+^j$ conditional on $\sum_{i=1}^n X_+^i = \eta$ has a binomial distribution with parameters $\eta$ and

$$\frac{v_j \int_0^{t_j} E_+(s)ds}{\sum_{i=1}^n v_i \int_0^{t_i} E_+(s)ds},$$

representing a standard property of independent Poisson random variables. It follows that the expected fraction of positively selected non-carcinogenic mutations over the whole cohort that belong to patient $j$ is

$$\alpha_j := \mathbb{E}\left[\frac{X_+^j}{\sum_{i=1}^n X_+^i}\middle|\sum_{i=1}^n X_+^i > 0\right] = \frac{v_j \int_0^{t_j} E_+(s)ds}{\sum_{i=1}^n v_i \int_0^{t_i} E_+(s)ds}.$$

Similarly, the expected fraction of neutral mutations who belong to patient $j$ is

$$\beta_j := \mathbb{E}\left[\frac{X_0^j}{\sum_{i=1}^n X_0^i}\middle|\sum_{i=1}^n X_0^i > 0\right] = \frac{v_j t_j}{\sum_{i=1}^n v_i t_i}.$$

Without loss of generality, say that $t_j$ is non-decreasing in $j$, and observe that

$$\frac{\alpha_j}{\beta_j} = t_j^{-1}\int_0^{t_j} E_+(s)ds \times \frac{\sum_{i=1}^n v_i t_i}{\sum_{i=1}^n v_i \int_0^{t_i} E_+(s)ds}$$

is non-decreasing in $j$ (which can be seen by taking the derivative of $t^{-1}\int_0^t E_+(s)ds$ with respect to $t$). Then by Lemma 3.2, for all $j$,

$$\mathbb{E}\left[\frac{\sum_{i=j}^n X_+^i}{\sum_{i=1}^n X_+^i}\middle|\sum_{i=1}^n X_+^i > 0\right] \geq \mathbb{E}\left[\frac{\sum_{i=j}^n X_0^i}{\sum_{i=1}^n X_0^i}\middle|\sum_{i=1}^n X_0^i > 0\right].$$

which implies the statement of the proposition. □

##### 3.2 Numerical calculations for Figures 4 and S4

Here we present the calculations for the null model in Figures 4 and S4. Say that mutation z arrives as a Poisson process $(K_t : t \geq 0)$ with constant rate $v$. Let $\tau_i = \min\{t \geq 0 : K_t = i\}$ denote the time of the $i$th arrival of z. Let $Y_i(t)$ be i.i.d. stochastic processes on the non-negative integers to represent the sizes of mutant-z clones at time $t$ after their initiation. Write $y(t) = \mathbb{E}Y_1(t)$ for the expected clone size. In Figure S4, the exponential model is $y(t) = e^{st}$, Gompertz $y(t) = e^{\log(1000)(1-e^{-st/\log(1000)})}$, logistic $y(t) = 1000/(1 + 999e^{-\frac{1000}{999}st})$, and polynomial $y(t) = (1 + st/3)^3$, where $s = \frac{dy}{dt}(0)$ is the growth rate parameter. The total number of mutant-z cells at time $t$ is

$$\sum_{i=1}^{K_t} Y_i(t-\tau_i),$$

whose expectation is

$$v\int_0^t y(u)du.$$

On the other hand, a neutral mutation $z_0$ with the same arrival rate has expected frequency $vt$. We model that cancer arrives at an age-dependent rate $r(t) = Ct^3$ for $t \in (0, 80)$, and that cancer arises independently of mutations $z$ and $z_0$. Over the population, cancer with mutation z arises at rate proportional to

$$r(t)\int_0^t y(u)du.$$

The expected frequency of mutation $z$ divided by the expected frequency of mutation $z_0$ among cancers is

$$\frac{\int_0^{80} r(t)\int_0^t y(u)dudt}{\int_0^{80} r(t)tdt},$$

and the mean age for mutation $z$ minus the mean age for mutation $z_0$ among cancers is

$$\frac{\int_0^{80} tr(t)\int_0^t y(u)dudt}{\int_0^{80} r(t)\int_0^t y(u)dudt} - \frac{\int_0^{80} t^2 r(t)dt}{\int_0^{80} tr(t)dt}.$$

### 4 Some branching process models of somatic evolution towards carcinogenesis

In this section we present some branching process models of somatic evolution to illustrate various musings on carcinogenic mutations' meaning and manifestation. The subsections can be read independently of each other, since they are conceptually and notationally distinct. Firstly, for the single mutation model of Figure 2, we point out mutation frequency differences between normal tissue in the absence of cancer, normal tissue conditional on cancer arrival, and cancer. Secondly, we extend the model to two interacting mutations, discussing how epistasis can confound comparison of cancer and normal tissue. We also see that mutational interactions can serve as a source of age-dependent carcinogenic effects. Thirdly, we extend the model further to multiple mutations, and in the special case of no epistasis, we observe agreement between a mutation's carcinogenic effect (1) and the observational hazard ratio for carcinogenesis given the mutation (2). Finally, extending/simplifying genetic space to infinite sites, we detail numerical calculations for Figure 6.

##### 4.1 A single mutation in cancerous and non-cancerous normal tissue

To begin, let's recall the model introduced in Figure 2, which focuses on a specific mutation named $\xi$. Let $\langle 1\rangle$ denote a cell with mutation $\xi$ and let $\langle 0\rangle$ denote a cell without the mutation. In a normal tissue, cells divide, die, mutate, and initiate cancer according to

$$\langle i\rangle \to \begin{cases} \langle i\rangle + \langle i\rangle, & \text{at rate } a_i \\ \emptyset, & \text{at rate } b_i \\ \langle 1\rangle, & \text{at rate } v \\ \text{cancer}, & \text{at rate } r_i \end{cases} \qquad \text{for } i \in \{0,1\},$$

with $a_0 = b_0$. We allow that the cancer rates $r_i$ vary with age. The mutation's carcinogenic effect (1) is $w = r_1/r_0$. In a tissue, or patch of tissue, at age $t$ let $M_1(t)$ and $M_0(t)$ respectively denote the number of non-cancerous cells with and without mutation $\xi$. Let $T$ denote the first time that cancer arrives (which would be infinite if cancer never arrives). Time $T$ is distributed as

$$\mathbb{P}[T > t] = \mathbb{E}e^{-\int_0^t r_1(s)M_1(s)+r_0(s)M_0(s)ds}.$$

Conditional on no cancer by time $t$, the expected numbers of cells with ($i = 1$) and without ($i = 0$) the mutation $\xi$ at time $t$ are

$$m_i(t) := \mathbb{E}\left[M_i(t)|T > t\right] = \frac{\mathbb{E}\left[M_i(t)e^{-\int_0^t r_1(s)M_1(s)+r_0(s)M_0(s)ds}\right]}{\mathbb{E}\left[e^{-\int_0^t r_1(s)M_1(s)+r_0(s)M_0(s)ds}\right]}.$$

By contrast, conditional on cancer before time $t$, the expected numbers of non-cancer cells with and without mutation $\xi$ at time $t$ are

$$m_i^*(t) := \mathbb{E}\left[M_i(t)|T < t\right] = \frac{\mathbb{E}\left[M_i(t)\left(1 - e^{-\int_0^t r_1(s)M_1(s)+r_0M_0(s)ds}\right)\right]}{\mathbb{E}\left[1 - e^{-\int_0^t r_1(s)M_1(s)+r_0(s)M_0(s)ds}\right]}.$$

So over a large cohort of tissues, by the law of large numbers, the odds of mutation $\xi$'s presence in a normal tissue cell sampled from non-cancerous (patches of) tissues at time $t$ is

$$\text{odds}_{\text{non-cancer}} = \frac{m_1(t)}{m_0(t)}$$

whereas the odds of mutation $\xi$'s presence in a normal tissue cell sampled from cancer-related lineages at time $t$ is

$$\text{odds}_{\text{cancer-related}} = \frac{m_1^*(t)}{m_0^*(t)}.$$

Let's compare these normal tissue mutation frequencies against cancers. Conditional on cancer arising for the first time at time $t$, the cancer founder cell carries mutation $\xi$ with odds

$$\text{odds}_{\text{cancer}} = \frac{\mathbb{E}\left[r_1(t)M_1(t)e^{-\int_0^t r_1(s)M_1(s)+r_0(s)M_0(s)ds}\right]}{\mathbb{E}\left[r_0(t)M_0(t)e^{-\int_0^t r_1(s)M_1(s)+r_0(s)M_0(s)ds}\right]} = w \times \text{odds}_{\text{non-cancer}};$$

indeed, in the language of (1) and (2), $Y$ plays no role, and so the carcinogenic effect and the observational hazard ratio for cancer given the mutation are trivially equivalent. However,

$$\text{odds}_{\text{cancer}} \neq w \times \text{odds}_{\text{cancer-related}}.$$

The basic conceptual point has been made: cancer-producing lineages in normal tissue provide a biased view of the frequencies of carcinogenic mutations in normal tissue, explaining why in our patient-matched comparison of cancerous and normal esophagus samples, we excluded 'normal' esophagus samples mutationally related to the cancers. Really, this point didn't need the specific branching process model to be made, but the model additionally serves the purpose of warming up for subsequent models of multiple mutations. We shall ask again about the relationship between mutation frequencies and the carcinogenic effect in the more interesting scenario of mutational interactions.

#### 4.2 Two epistatic mutations

We extend the model to consider interactions between two mutations, named 1 and 2. Here, $z_1 = 1$ and $z_1 = 0$ respectively represent that mutation 1 is present/absent in a cell, while $z_2 = 1$ and $z_2 = 0$ represent the same thing for mutation 2. A cell's genotype is $(z_1, z_2)$, an element of $\{0,1\}^2$. A cell with genotype $(i,j) \in \{0,1\}^2$ divides, dies, mutates, and transforms to cancer according to

$$(i,j) \to \begin{cases} (i,j) + (i,j), & \text{at rate } a_{i,j} \\ \emptyset, & \text{at rate } b_{i,j} \\ (1,j), & \text{at rate } v_1 \\ (i,1), & \text{at rate } v_2 \\ \text{cancer}, & \text{at rate } r_{i,j}, \end{cases}$$

with $a_{0,0} = b_{0,0}$. For this model, we shall examine the how the meaning and measurement of a mutation's carcinogenic effect may depend on another mutation.

Across a cohort of people, let $f_{i,j}(t)$ denote the fraction of normal tissue cells at age $t$ with genotype $(i,j)$. Artificially perturbing the status of mutation 1, we see that the carcinogenic effect (1) of mutation 1 is

$$\frac{(f_{0,0}(t) + f_{1,0}(t))r_{1,0} + (f_{0,1}(t) + f_{1,1}(t))r_{1,1}}{(f_{0,0}(t) + f_{1,0}(t))r_{0,0} + (f_{0,1}(t) + f_{1,1}(t))r_{0,1}}. \tag{5}$$

For a sanity check, consider the special case $r_{1,1}/r_{0,1} = r_{1,0}/r_{0,0} = w$, meaning that mutation 1's effect does not depend upon mutation 2: reassuringly, (5) simplifies to $w$. For a slightly more interesting special case, consider $r_{0,0} = r_{0,1} = r_{1,0}$ and write $\omega = r_{1,1}/r_{0,0}$ for the effect of the two mutations together, so (5) simplifies to

$$f_{0,0}(t) + f_{1,0}(t) + \omega(f_{0,1}(t) + f_{1,1}(t));$$

similarly, the carcinogenic effect of mutation 2 is $f_{0,0}(t)+f_{0,1}(t)+\omega(f_{1,0}(t)+f_{1,1}(t))$. This second special case is instructive: although the mutations equally need each other to affect carcinogenesis, they may still have distinct carcinogenic effects due to their differing frequencies in normal tissue: mutation 1's carcinogenicity depends upon mutation 2's prevalence, and vice versa.

We return to the more general case of mutation 1's carcinogenic effect (5). For comparison, the observational hazard ratio (2) for cancerous transformation given mutation 1 is

$$\frac{\frac{f_{1,0}(t)}{f_{1,0}(t)+f_{1,1}(t)} r_{1,0} + \frac{f_{1,1}(t)}{f_{1,0}(t)+f_{1,1}(t)} r_{1,1}}{\frac{f_{0,0}(t)}{f_{0,0}(t)+f_{0,1}(t)} r_{0,0} + \frac{f_{0,1}(t)}{f_{0,0}(t)+f_{0,1}(t)} r_{0,1}}. \tag{6}$$

Mutation 1's carcinogenic effect (5) and observational hazard ratio (6) are equal under the condition

$$f_{i,j}(t) = (f_{i,0}(t) + f_{i,1}(t))(f_{0,j}(t) + f_{1,j}(t)), \tag{7}$$

which says that mutations 1 and 2 are independently distributed among normal tissue cells.

We ask: What somatic evolutionary parameters provide condition (7)? To answer, we seek an expression for the $f_{i,j}(t)$. To begin, observe the Kolmogorov equations

$$\frac{d}{dt} n_{i,j} = (\gamma_{i,j} - r_{i,j}) n_{i,j} + (-1)^{i+1} v_1 n_{0,j} + (-1)^{j+1} v_2 n_{i,0}$$

for the expected number of normal cells $n_{i,j}(t)$ at age $t$ with genotype $(i,j)$, where $\gamma_{i,j} = a_{i,j} - b_{i,j}$ denotes the fitness of genotype $(i,j)$. For analytic tractability, we focus on the limit of small $v_1, v_2, r_{i,j}$, so we don't worry about conditioning on whether or not cancer has arrived like we did for the single mutation model. Given initial condition $n_{0,0}(0) = n$ and $n_{i,j}(0) = 0$ for $i + j > 0$, the solution of the Kolmogorov equations is

$$\begin{cases} n_{0,0}(t) = n \\ n_{1,0}(t) = \frac{nv_1}{\gamma_{1,0}}(e^{\gamma_{1,0}t} - 1) \\ n_{0,1}(t) = \frac{nv_2}{\gamma_{0,1}}(e^{\gamma_{0,1}t} - 1) \\ n_{1,1}(t) = c_{1,1}e^{\gamma_{1,1}t} + c_{1,0}e^{\gamma_{1,0}t} + c_{0,1}e^{\gamma_{0,1}t} + c_{0,0}, \end{cases}$$

where $c_{0,0} = \frac{nv_1v_2(\gamma_{1,0}+\gamma_{0,1})}{\gamma_{0,1}\gamma_{1,0}\gamma_{1,1}}$, $c_{1,0} = -\frac{nv_1v_2}{\gamma_{1,0}(\gamma_{1,1}-\gamma_{1,0})}$, $c_{0,1} = -\frac{nv_1v_2}{\gamma_{0,1}(\gamma_{1,1}-\gamma_{0,1})}$, and $c_{1,1} = -c_{0,0} - c_{1,0} - c_{0,1}$. Then over an infinite cohort of people at age $t$, by the law of large numbers, the fraction of normal cells with genotype $(i,j)$ is $f_{i,j}(t) = n_{i,j}(t)/(n_{0,0}(t) + n_{1,0}(t) + n_{0,1}(t) + n_{1,1}(t))$, which in our parameter regime of small $v_i$ simplifies to $f_{i,j}(t) = n_{i,j}(t)/n$. Here, we observe that condition (7) is equivalent to

$$c_{1,1}e^{\gamma_{1,1}t} + c_{1,0}e^{\gamma_{1,0}t} + c_{0,1}e^{\gamma_{0,1}t} + c_{0,0} = \frac{nv_1v_2}{\gamma_{1,0}\gamma_{0,1}}(e^{\gamma_{1,0}t} - 1)(e^{\gamma_{0,1}t} - 1),$$

which holds if and only if $\gamma_{1,1} = \gamma_{1,0} + \gamma_{0,1}$, i.e. the mutations' selective effects in normal tissue are additive. We conclude that epistasis during normal tissue evolution is a potential source of disagreement between the carcinogenic effect and observational hazard ratio.

Next, focussing on the zero epistasis case, we extend the two-mutation model to allow multiple mutations, acting as a bridge towards our infinite-sites model that will follow.

#### 4.3 Multiple independent mutations

We extend genetic space to the set $\mathcal{G} = \{0,1\}^L$ of binary sequences of arbitary length $L$. That is, a cell's genotype is depicted by a finite sequence where each entry of the sequence represents a genomic position, and a position takes the value zero or one to denote that it is unmutated or mutated respectively. Initially there are $n$ cells with genotype $(0,..,0)$. Position $i$ mutates at rate $v_i$ per cell: along a single lineage, each genomic position independently follows a continuous-time Markov process on $\{0,1\}$ that begins at 0 and transitions to 1 at rate $v_i$. A cell with genotype $z = (z_1,..,z_L) \in \mathcal{G}$ divides at rate $a(z)$, is lost through death or differentiation at rate $b(z)$, or transforms to cancer at rate $r(z)$. In other words,

$$\langle z\rangle \to \begin{cases} \langle z\rangle + \langle z\rangle, & \text{at rate } a(z) \\ \emptyset, & \text{at rate } b(z) \\ \langle z[z_i = 1]\rangle, & \text{at rate } v_i \\ \text{cancer}, & \text{at rate } r(z), \end{cases}$$

where $\langle z\rangle$ represents a cell with genotype $z$ and $\langle z[z_i = 1]\rangle$ represents a cell whose mutations agree with $z$ except possibly at position $i$ which is set to 1.

Given the exponential grandiosity of parameter space, we impose the restrictive condition

$$a(z) - b(z) = \sum_{i=1}^{L} z_i s_i$$

of zero epistasis in normal tissue, where $s_i \in \mathbb{R}$ is the selective effect in normal tissue of a mutation at position $i$, and the condition

$$r(z) = r(0)\prod_{i=1}^{L} w_i^{z_i}$$

of multiplicative mutational interactions at carcinogenesis, where $w_i \in (0,\infty)$ is the carcinogenic effect of a mutation at position $i$.

For a genotype $z$, write $z[z_i = 0]$ for the genotype whose mutations agree exactly with those of $z$ except possibly at position $i$ where $z[z_i = 0]$ has no mutation. Then the expected number of normal cells with genotype $z$ at time $t$, $n_z(t)$, follows the forward Kolmogorov equations

$$\frac{dn_z}{dt} = n_z\left(\sum_{i=1}^{L}(z_i s_i - (1-z_i)v_i) - r(0)\prod_{i=1}^{L} w_i^{z_i}\right) + \sum_{i=1}^{L} n_{z[z_i=0]} z_i v_i \tag{8}$$

with initial condition

$$n_z(0) = \begin{cases} n, & \text{for } z = (0,..,0) \\ 0, & \text{otherwise.} \end{cases}$$

The solution in the limit of small $v_i$ and $r(0)$ is

$$n_z = n \prod_{i=1}^{L} \left( v_i \frac{e^{s_i t} - 1}{s_i} \right)^{z_i}. \tag{9}$$

We find that the observational hazard ratio (2) for cancerous transformation given mutation $i$

$$\frac{\sum_{z \in \mathcal{G}: z_i = 1} n_z(t) r(z) / \sum_{z \in \mathcal{G}: z_i = 1} n_z(t)}{\sum_{z \in \mathcal{G}: z_i = 0} n_z(t) r(z) / \sum_{z \in \mathcal{G}: z_i = 0} n_z(t)}$$

is equal to the carcinogenic effect of mutation $i$

$$\frac{\sum_{z \in \mathcal{G}} n_z(t) r(z[z_i = 1])}{\sum_{z \in \mathcal{G}} n_z(t) r(z[z_i = 0])} = w_i.$$

This result might look unsurprising; its banality is reassuring. There are of course other important statistics too – on the relationships between patient age, number of carcinogenic mutations per cancer, and the mutations' carcinogenic effects – whose calculation we defer to a refined model of genetic evolution.

#### 4.4 Infinite sites; numerical calculations for Figures 6 and S6

Moving forward from the just-posed multiple mutation model, we refine genetic space to infinite dimensions, corresponding to the limit $L \to \infty$ and $v_i \downarrow 0$ with the product $v_i L > 0$ fixed. The virtue of an infinite genome is that it simplifies parameter space while allowing a continuous spectrum for the effects of mutations. Here, it is mutations' carcinogenic effects rather than selective effects that we focus upon. For simplicity, we forget about cell divisions and deaths in normal tissue. We study the acquisition of mutations along a single lineage in normal tissue. Let $K = (K_t : 0 \le t \le T)$ be a rate $v$ Poisson process denoting the acquisition of mutations. Let $W = (W_1, W_2, ..)$ be an i.i.d. sequence of non-negative random variables denoting the mutations' carcinogenic effects. Let $Z = (Z_t : 0 \le t \le T)$ be such that conditional on $K$ and $W$, $Z$ is a continuous-time Markov process on $\{0, 1\}$ with initial condition $Z_0 = 0$ and with a transition from 0 to 1 at rate $r_0 \prod_{i=1}^{K_t} W_i$ at time $t$. The cancer initiation time is $\tau = \min\{t \ge 0 : Z_t = 1\}$. This model can be regarded as an adaptation of Armitage and Doll's, and Nordling's, classic multi-stage model of carcinogenesis. The key difference is that we allow mutations to vary in their cancer-causing strength.

Now we detail the numerical recipe underlying Figures 6 and S6. We begin with Figure S6A. Write $r_k = r_0 \prod_{i=1}^{k} w_i$ for the rate at which a cell initiates cancer conditional on $k$ mutations with carcinogenic effects $w_1, .., w_k$. Observe that $g_k(t; w) = \mathbb{P}[K_t = k, Z_t = 0 | W = w]$ follows

$$\frac{dg_k}{dt} = \begin{cases} -v g_0, & k = 0 \\ v g_{k-1} - (v + r_k)\, g_k, & k \ge 1 \end{cases} \tag{10}$$

with initial condition $g_k(0; w) = \delta_{0,k}$. The solution is

$$g_k(t; w) = \sum_{j=0}^{k} \frac{(-v)^k e^{-(v + r_j)t}}{\prod_{\substack{i=1 \\ i \ne j}}^{k} (r_j - r_i)},$$

or alternatively put into the computer the matrix exponential solution of (10). Calculate $g_k(t; w) = \mathbb{P}[K_t = k, Z_t = 0 | W = w]$ for each of a bunch of independent simulations of $W$ – call them $w_{(1)}, .., w_{(R)}$ – and use the approximation

$$\mathbb{P}[K_t = k, Z_t = 0] \approx \frac{1}{R} \sum_{r=1}^{R} \mathbb{P}[K_t = k, Z_t = 0 | W = w_{(r)}],$$

from which we obtain the probability distribution of mutation burden among normal tissue cells

$$\mathbb{P}[K_t = k | Z_t = 0] = \frac{\mathbb{P}[K_t = k, Z_t = 0]}{\sum_j \mathbb{P}[K_t = j, Z_t = 0]}.$$

That is part of Figure S6A. The other part is the probability distribution of mutation burden among cancer founders, for which we will again use the conditional distribution of mutation numbers in normal cells $g_k(t) = \mathbb{P}[K_t = k, Z_t = 0|W = w]$ calculated as before. Observe that

$$\mathbb{P}[K_t = k, \tau \in (t, t + dt)|W = w] = \mathbb{P}[K_t = k, Z_t = 0|W = w] \times r_0 \prod_{i=1}^{k} w_i \times dt. \tag{11}$$

As before, average over simulated $w$, and normalise, to obtain $\mathbb{P}[K_t = k|\tau = t]$. That concludes Figure S6A.

Next towards Figure 6B, simulate a load of independent $W_i$, named $w_{i,r}$, and use (11) to calculate

$$\begin{aligned} &\mathbb{P}[K_t = k, \tau \in (t, t + dt), W_j \in (w + dw)] \qquad (12)\\ &\approx \frac{1}{R}\sum_{r=1}^{R} \mathbb{P}[K_t = k, \tau \in (t, t + dt)|W_j = w \text{ and } W_i = w_{i,r} \text{ for } i \in \{1, .., k\} - \{j\}] \times f(w)dw, \end{aligned}$$

where $f$ is the probability density for $W_i$. Then

$$\mathbb{P}[W_j \in (w + dw)|K_\tau = k] = \frac{\int \mathbb{P}[K_t = k, \tau \in (t, t + dt), W_j \in (w + dw)]dt}{\int\int \mathbb{P}[K_t = k, \tau \in (t, t + dt), W_j \in (w + dw)]dwdt}.$$

And finally

$$\mathbb{E}\left[K_\tau^{-1} \sum_{i=1}^{K_\tau} I(W_i \in (w, w + dw))\big|K_\tau = k\right] = k^{-1} \sum_{i=1}^{k} \mathbb{P}\left[W_i \in (w, w + dw)|K_\tau = k\right]$$

gives 6B. For 6C, revisit (12) to get

$$\begin{aligned} &\mathbb{E}\left[K_\tau^{-1} \sum_{i=1}^{K_\tau} I(W_i \in (w, w + dw))\big|\tau = t, K_\tau > 0\right] \\ &= \frac{\sum_{k\geq 1} k^{-1} \sum_{i=1}^{k} \mathbb{P}[K_t = k, \tau \in (t, t + dt), W_i \in (w + dw)]}{\mathbb{P}[\tau \in (t, t + dt), K_\tau > 0]}. \end{aligned}$$

### 5 Criticisms and alternatives for the carcinogenic effect definition

Since the carcinogenic effect of a mutation is a key definition in our paper, we want to attack our definition, to think about whether it makes sense.

#### 5.1 Causation is imaginary

We must ask: is our notion of cancer causation reasonable or nonsensical? First we turn to the more general and foundational question: is *any* notion of causation reasonable or are they all nonsensical?

Bertrand Russell, one of the most respected philosophers of the 20th century, provided a devastating critique of causation[1]. He wrote "*All philosophers, of every school, imagine that causation is one of the fundamental axioms or postulates of science, yet, oddly enough, in advanced sciences such as gravitational astronomy, the word 'cause' never occurs. .. To me it seems .. that the reason why physics has ceased to look for causes is that, in fact, there are no such things. The law of causality, I believe, like much that passes muster among philosophers, is a relic of a bygone age, surviving, like the monarchy, only because it is erroneously supposed to do no harm.*" Russell proceeded to logically demolish dictionary definitions of causation that were in common usage, declaring the definitions to be nonsense. That was in 1912. Now, more than a century later, standard modern day dictionaries offer definitions of causation that do not appear to differ from those that Russell earlier destroyed.

Modern day biologists and other scientists have a more powerful notion of causation than any standard dictionary. Biologists measure causation as the statistical outcome of a controlled experiment. For example,

[1]Bertrand Russell, On the Notion of Cause, Proceedings of the Aristotelian Society, 1912–1913

suppose that one group of mice were given coca cola to drink while another group of mice were given water. If there were a statistically significant elevation in cancer incidence among the coca cola- relative to water-drinking mice, then the biologists performing the experiment would say that coca cola likely causes cancer in mice. Elegantly, this experimental and statistical framework allows assessment of causation regardless of ignorance of the many variables besides coca cola that may contribute to cancer. However, definitive determination of coca cola's causal contribution is precluded by statistical noise. Statistical noise can only be calmed by an *infinite* number of mice. In general, an infinite-sized controlled experiment can estimate x's causal effect on y with perfect precision. An infinite-sized controlled experiment can therefore be regarded as the modern day biological meaning of causation.

Thus, biological causation is like a Platonic ideal. It has as much basis in the physical universe as frequentist probability. Causation does have a physical approximation that is a finite controlled experiment. Sometimes though, causation is claimed even where no controlled experiment, even a finite one, is physically performed. Economists commonly use the word causation in the absence of controlled experiments, instead appealing to natural experiments for a mimicry of the controlled. Biologists likewise talk of causes where the relevant controlled experiment is impossible. For instance, cancer geneticists have declared that *TP53* mutations cause human cancers. Cancer geneticists think that artificial insertion of *TP53* mutations into human tissues in vivo would lead to elevated rates of cancer, even though this definitional experiment can never be enacted. Instead, their conviction that *TP53* mutations cause human cancer comes from countless lines of indirect evidence of causation, including human observational studies and mouse genetic-editing experiments who are imperfect proxies for the hypothetical human genetic-editing experiment. The indirect evidence further includes studies of *mechanism*. Abstractly, a mechanistic explanation for how x causes y is a sequence of small causal arrows who together form a large causal arrow from x to y. One small causal arrow points from a *TP53* mutation to impaired p53 production, another small causal arrow points from impaired p53 production to faulty apoptosis, and so on. Again however, these causal arrows lack measurement by controlled experiments in human tissues in vivo. Despite widespread belief in the mutational causation of human cancer, the strict experimental definition for causation does not take place in our observable physical universe – belief takes place in our minds.

This is not a dismissal of causation. Imaginary is not equivalent to nonsensical. Would Russell condemn the hypotheses that *TP53* mutations and sugary drinks cause human cancers? Would he say that they are nonsense? No. He would respect biologists' concept of causation in the same way as he would respect mathematicians' concept of a circle or their axioms of set theory – without any expectation of perfect physical embodiment, but with the demand for precise definitions.

#### 5.2 Selecting a suitable experiment

We define mutational causation of cancer in terms of an imaginary genetic editing experiment. In brief, we genetically edit cells of a specific type in vivo among an infinite number of people: we randomly assign mutation $\xi$ to some cells while disallowing mutation $\xi$ from the other cells (the fraction of cells with the mutation varying between people). Then, the per cell hazard ratio for cancer initiation given mutation $\xi$ defines the mutation's carcinogenic effect. This experiment is just one of many possible experiments that could reasonably define mutational causation of cancer. So, why did we choose this one? For a comparative discussion, let's name our carcinogenic effect definition as *Definition 0*, and now here are three alternatives:

*Definition 1*: Partition an infinite cohort of people into two groups. Introduce mutation $\xi$ to all cells of each person of one group, and disallow mutation $\xi$ from the other group. The carcinogenic effect is the per person hazard ratio for cancer diagnoses given the mutation.

*Definition 2*: Partition an infinite cohort of people into two groups. Introduce mutation $\xi$ to one cell per person of one group but not the other group. The carcinogenic effect is a person's lifetime odds ratio for a cancer diagnosis in the mutation vs non-mutation group.

*Definition 3*: Partition an infinite cohort of people into two groups. Leave one group of people untouched (so mutation $\xi$'s frequency in normal tissue follows natural somatic evolution) and permanently disallow mutation $\xi$ from the other group. The carcinogenic effect is [one minus lifetime risk of cancer in the non-mutation group] divided by [lifetime risk of cancer in the mutation group], i.e. the proportion of lifetime cancer risk explained by mutation $\xi$.

Besides Definitions 0, 1, 2, and 3, of course other definitions are possible too. Our purpose here is not to provide an exhaustive list. It is to show that multiple options exist, to provide material for criticism of Definition 0, and to explain why we settled upon Definition 0 amongst other options.

To begin nitpicking, we note something unusual about Definition 0's experiment: the experimental unit is a cell, and cells can divide through the experiment. In Definitions 1, 2 and 3, by contrast, the experimental unit is a person, and no person can divide themself into two near-identical people. In real-world experiments too across science, experimental units do not replicate. Definition 0 thus deviates from the standard experimental format. Breaking tradition per se is no concern. We are not slaves to tradition. Nevertheless, we prefer people over cells as the experimental unit, for two reasons, both relating to cell divisions. Firstly, while cancer development is naturally modelled as independent between people, cancer development is not so naturally modelled as independent between cells – cell divisions introduce statistical dependencies between cells. Dependencies may also arise due to physical space or cell-cell signalling within a tissue. Lack of independence between cells within a tissue is not an indictment of Definition 0, but it is an inelegance. The second reason we prefer people over cells as the experimental unit is that for people, the experimental measurement can be cancer diagnosis, whereas for cells, the experimental measurement is the transformation from normal cell to cancer cell. The advantage of the former is that cancer diagnosis is an unambiguous statement provided by a doctor; the disadvantage of the latter is that the normal-cancer transformation at the level of the single cell might not be an unambiguous point in time, unless we provide an auxiliary definition for the point of cancer transformation, such as the birth of the most recent common ancestor of all cells who comprise a cancer diagnosed by a doctor.

Another criterion by which to compare the carcinogenic effect definitions is their depictions of the natural distribution of mutations in human tissues. Definition 1's experiment, by enforcing genetic homogeneity, could be a reasonable representative for germline mutations but it could not be a reasonable representative for somatic mutations. Definition 2's experiment, by introducing only a single mutation, could be a reasonable representative for somatic mutations, but only for those mutations who arise extremely rarely in a tissue. Definition 3's experiment's control group does not have an analogue in real human biology that I am aware of. Definition 0's experiment on the other hand, allows an arbitrary fraction of cells per person with mutation $\xi$, as does somatic evolution, which is a major point in its favour.

We like Definition 0 furthermore for its clean distinction between cancer causation and selection in normal tissue. Neither does Definition 1 have any relationship with selection in normal tissue (in fact, Definitions 0 and 1 are equivalent under the assumptions that cells act independently of each other, that the number of normal cells per person is independent of whether the person is given the mutation, and that the time between cancer initiation and diagnosis can be neglected). Definitions 2 and 3 offer slightly different yardsticks which do depend upon the mutation's selective effect in normal tissue – a greater selective effect in normal tissue implies a greater mutation frequency in normal tissue which, for a carcinogenic mutation, implies a greater cancer risk and a therefore a greater cancer-causing effect by these definitions. Definition 3 additionally depends upon the rate of mutation acquisition. This is not a criticism of Definitions 2 and 3 but an expression of subjective preference for Definition 0. Distinguishing selection and causation is a major purpose of our paper, so we like to define a mutation's carcinogenic effect orthogonally to the selective effect in normal tissue. Relatedly, Definition 0 hones in on what makes cancer genomes special relative to normal tissue: the greater a mutation's carcinogenic effect, the more that the mutation distinguishes cancer from normal tissue genomes, with therapeutic implications.
